# Translation is a key determinant controlling the fate of cytoplasmic long non-coding RNAs

**DOI:** 10.1101/2022.05.25.493276

**Authors:** Sara Andjus, Ugo Szachnowski, Nicolas Vogt, Isabelle Hatin, David Cornu, Chris Papadopoulos, Anne Lopes, Olivier Namy, Maxime Wery, Antonin Morillon

**Affiliations:** ncRNA, epigenetic and genome fluidity, Institut Curie, PSL University, Sorbonne Université, CNRS UMR3244, 26 rue d’Ulm, F-75248 Paris Cedex 05, France; ncRNA, epigenetic and genome fluidity, Institut Curie, Sorbonne Université, CNRS UMR3244, 26 rue d’Ulm, F-75248 Paris Cedex 05, France; Genomics, Structure and Translation, Institute for Integrative Biology of the Cell (I2BC), CEA, CNRS, Université Paris-Sud, Université Paris-Saclay, 91198 Gif-sur-Yvette cedex, France; Molecular Bio-informatics, Institute for Integrative Biology of the Cell (I2BC), CEA, CNRS, Université Paris-Sud, Université Paris-Saclay, 91198 Gif-sur-Yvette cedex, France

**Author notes:** Co-first authors (Sara Andjus, Ugo Szachnowski). Corresponding & co-last authors.

**Keywords:** lncRNA, Xrn1, NMD, translation

## Abstract

Despite being predicted to lack coding potential, cytoplasmic long non-coding (lnc)RNAs can associate with ribosomes, which may result in the production of functional peptides. However, the landscape and biological relevance of lncRNAs translation remains poorly studied. In the budding yeast *Saccharomyces cerevisiae*, cytoplasmic Xrn1-sensitive lncRNAs (XUTs) are targeted by the Nonsense-Mediated mRNA Decay (NMD), suggesting a translation-dependent degradation process. Here, we report that XUTs are translated, which impacts their abundance. We show that XUTs globally accumulate upon translation elongation inhibition, but not when initial ribosome loading is impaired. Translation also affects XUTs independently of NMD, in some cases interfering with their decapping. Ribo-Seq confirmed ribosomes binding to XUTs and identified actively translated small ORFs in their 5’-proximal region. Mechanistic analyses revealed that their NMD-sensitivity mainly depends on the 3’-untranslated region length. Finally, we detected the peptide derived from the translation of an NMD-sensitive XUT reporter in NMD-competent cells. Our work highlights the role of translation in the post-transcriptional metabolism of XUTs, acting as a modulator of their expression. We propose that XUT-derived peptides could be exposed to the natural selection, while NMD restricts XUTs levels.

## INTRODUCTION

Long non-coding (lnc)RNAs constitute a class of transcripts that arise from the pervasive transcription of eukaryotic genomes (1). Although the debate on their functional significance is still ongoing (2), some of them are now recognized as important RNA regulators involved in multiple cellular functions (3–5). Consistent with their functional importance, their expression appears to be precisely controlled (6,7). Furthermore, the abnormal expression of lncRNAs is associated with human diseases, including cancers (8–10). However, such evidence remains marginal and a full mechanistic description is still required to understand the *raison d’être* of lncRNAs in cells, as well as the molecular mechanisms regulating their expression.

By definition, lncRNAs have been predicted to lack coding potential. However, this assumption has been challenged by several independent observations, which have shown that cytoplasmic lncRNAs can associate with ribosomes (11–15). In fact, ribosome profiling (Ribo-Seq) analyses identified small open reading frames (smORFs) on subsets of lncRNAs (11,15–18), and in some cases, their translation even results into the production of functional peptides (19–23).

Apart from such examples of functional lncRNA-derived peptides, which are marginal to date, the extent and the biological relevance of lncRNA translation remains unclear. An emerging view in the field proposes that lncRNAs could constitute a reservoir of rapidly evolving smORFs which the cell could exploit as a source of potential genetic novelty by producing novel peptides (24). If beneficial, lncRNA-derived peptides could be selected, thereby contributing to the emergence of novel protein-coding genes through the evolutionary process known as *de novo* gene birth (25–31).

In the budding yeast *Saccharomyces cerevisiae*, the idea that cytoplasmic lncRNAs can be translated has been suggested by their sensitivity to the Nonsense-Mediated mRNA Decay (NMD). NMD is a conserved translation-dependent RNA decay pathway known to target mRNAs bearing premature stop codons (32) as well as ‘normal’ mRNAs with a long 3’ untranslated region (UTR) (33–35); nevertheless, such ‘aberrant’ transcripts represent only one type of NMD substrates (see (36) for review). In fact, 70% of yeast cytoplasmic lncRNAs, known as Xrn1-sensitive Unstable Transcripts (XUTs) due to their extensive degradation by the cytoplasmic 5’-exonuclease Xrn1 (37), are NMD substrates (38,39). However, direct experimental evidence that XUTs are globally translated is still missing. Whether NMD-resistant lncRNAs (30% of XUTs) are also translated, and if so, what are the molecular features allowing them to escape NMD, remain open questions. The molecular consequences of translation on the post-transcriptional regulation of XUTs expression are also far from being understood. Finally, the output and biological relevance of such non-canonical translation events are still largely unknown.

Here, we investigated the impact of translation on the fate of cytoplasmic lncRNAs, using XUTs as a paradigm. We found that most NMD-sensitive and NMD-resistant XUTs rapidly accumulate in wild-type (WT) yeast cells treated with translation elongation inhibitors. Besides NMD, our data indicate that translation can also affect XUTs decay in an NMD-independent manner, by interfering with their decapping. In contrast to the effect of the translation elongation inhibitors, we found that XUTs levels remain unchanged in stress conditions associated with global inhibition of translation initiation, suggesting a mechanism where the elongating ribosomes protect them from the decay factors while they are translated. Ribo-Seq analyses confirmed that both NMD-sensitive and -resistant XUTs are actually bound by ribosomes, and we identified actively translated smORFs in their 5’ proximal portion. Mechanistic analyses on a candidate XUT demonstrated that its NMD-sensitivity depends on the length of its 3’ UTR. Finally, we show that peptides can be produced from an NMD-sensitive lncRNA reporter in WT cells, suggesting that despite the ‘cryptic’ nature of the transcript, its translation can result into a detectable product.

Altogether, our data support a model where translation occupies a central role in the metabolism of cytoplasmic lncRNAs, a rapid binding by ribosomes probably being the default route as they reach the cytoplasm. We propose that these translation events could allow lncRNA-derived peptides to be exposed to the natural selection, while NMD ensures that the majority of the transcripts they originate from are rapidly eliminated.

## MATERIAL AND METHODS

### Yeast strains and media

The strains used in this study are listed in Table S3. Mutants were constructed by transformation and were all verified by PCR on genomic DNA (see above).

Yeast cells were grown to mid-log phase (OD_600_ 0.5) at 30°C in Yeast Extract-Peptone-Dextrose (YPD) medium or Complete Synthetic Medium (CSM), with 2% glucose. In the glucose starvation experiments, glucose was replaced glycerol and ethanol.

5-Fluoroorotic acid (5-FOA) was used at a final concentration of 1 g/L on solid CSM plates. G418 (Geneticin; Gibco) was used at a final concentration of 100 µg/ml on solid YPD plates. CHX (Sigma) and ANS (Sigma) were used at a final concentration of 100 µg/ml.

### Construction of xut0741 mutants

The *xut0741-a*, *-b*, *-d* and *-f* alleles, flanked by NaeI sites, were produced as synthetic gBlocks DNA fragments (IDT – Integrated DNA Technologies), and then cloned between the KpnI and XbaI sites of the pAM376 backbone vector (40), giving the pAM594, pAM596, pAM598 and pAM600 vectors, respectively. The *xut0741-c* and *xut0741-e* mutants were constructed by site-directed mutagenesis from *xut0741-d* and then cloned into the same backbone vector, giving the pAM724 and pAM723 vectors, respectively. The sequence of each alleles was verified by Sanger sequencing and is available in Supplemental File 1. The mutant alleles were excised from the pCRII vector using NaeI digestion and transformed into the YAM2831 (where the *XUT0741*/*ADH2* locus has been deleted by *URA3*). After 1 day of growth on non-selective medium, transformants were replicated on CSM + 5-FOA plates and incubated at 30°C for 4-5 days. The proper integration of the mutant alleles was confirmed by PCR on genomic DNA using oligonucleotide AMO3350-3351. *UPF1* was deleted subsequently by transformation with the product of a PCR on YAM202 (*upf1Δ::kanMX4*) genomic DNA with oligonucleotides AMO2710-2711. The transformants were selected on YPD + G418 plates and *UPF1* deletion was verified by PCR on genomic DNA using oligonucleotides AMO190-2712.

The chimera-encoding plasmid (pAM726) was produced in two steps. Firstly, the 3’ UTR of the native *XUT0741* was amplified by PCR on YAM1 genomic DNA using oligonucleotides AMO3471-3382, and then cloned between the KpnI and XbaI sites of a pCRII-TOPO backbone, giving the pAM725 vector. Secondly, the sequence corresponding to the 5’ UTR and ORF of the *xut0741-d* mutant was amplified by PCR on YAM2854 genomic DNA using oligonucleotides AMO3379-3497, and then cloned between the KpnI and EcoRI sites of pAM725, giving the pAM726 vector. The sequence of the chimera allele was verified by Sanger sequencing (see Supplemental File 1). Plasmid digestion, transformation in YAM2831 cells, transformants selection and screening, as well as *UPF1* deletion, were as described above.

C-terminal 3FLAG tagging of *xut0741-b* was performed using an ‘overlap extension PCR’ strategy. A first amplicon was produced by PCR on YAM2853 genomic DNA using oligonucleotides AMO3379-3530. A second amplicon was produced by PCR on the same DNA using oligonucleotides AMO3382-3531. After purification on agarose gel, the two amplicons (displaying a 28-bp overlap) were mixed and used as DNA templates for PCR using oligonucleotides AMO3379-3382. The final full PCR product was then digested by KpnI and XbaI and cloned in the same backbone vector as the other *xut0741* mutants, giving the pAM728 plasmid (see Supplemental File 1 for insert sequence). All subsequent steps were as above.

The stem-loop (GATCCCGCGGTTCGCCGCGG), previously shown to inhibit *MFA2* mRNA translation (41), was inserted into the 3FLAG-tagged *xut0741-b* allele using a similar ‘overlap extension PCR’ strategy, involving the overlapping oligonucleotides AMO3550 (for the 5’ amplicon) and AMO3549 (for the 3’ amplicon), ultimately giving the pAM741 plasmid (insert sequence available in Supplemental File 1). All subsequent steps were as above.

*XRN1* was deleted in strains YAM2908 (*xut0741-b-3FLAG*) and YAM2934 (*SL-xut0741-b-3FLAG*) by transformation with the product of a PCR on YAM6 (*xrn1Δ::kanMX4*) genomic DNA with oligonucleotides AMO34-35. The transformants were selected on YPD + G418 plates and screened by PCR on genomic DNA using oligonucleotides AMO3247-1669.

### Total RNA extraction

Total RNA was extracted from exponentially growing cells (OD_600_ 0.5) using standard hot phenol procedure. Extracted RNA was ethanol-precipitated, resuspended in nuclease-free H_2_O (Ambion) and quantified using a NanoDrop 2000c spectrophotometer and/or a Qubit fluorometer with the Qubit RNA HS Assay kit (Life Technologies).

### Northern blot

10 μg of total RNA were separated on denaturing 1.2% agarose gel and then transferred to Hybond-XL nylon membrane (GE Healthcare). ^32^P-labelled oligonucleotides (listed in Table S4) were hybridized overnight at 42°C in ULTRAhyb®-Oligo hybridization buffer (Ambion). After hybridization, membranes were washed twice in 2X SSC/0.1% SDS for 15 minutes at 25°C, and once in 0.1X SSC/0.1% SDS for 15 minutes at 25°C. Membranes were exposed to Storage Phosphor screens. Signal was detected using a Typhoon Trio PhosphorImager and analysed with the version 10.1 of the ImageQuant TL sofware (Cytiva).

### Strand-specific RT-qPCR

Strand-specific RT-qPCR experiments were performed from three biological replicates, as previously described (42). The oligonucleotides used are listed in Table S4.

### Total RNA-Seq

For each strain/condition, total RNA-Seq was performed from two biological replicates. For each sample, 1 µg of total RNA was mixed with 2 µl of diluted ERCC RNA spike-in mix (1:100 dilution in nuclease-free H_2_O; Invitrogen). Ribosomal (r)RNAs were depleted using the Ribominus Eukaryote v2 kit (Ambion). Alternatively, 1.5 µg of total RNA was mixed with 3 µl of diluted ERCC RNA spike-in mix and then digested for 1h at 30°C with 1 unit of Terminator 5’-Phosphate-Dependent Exonuclease (Epicentre) in 1X Reaction Buffer A containing 10 units of SUPERase-In RNase inhibitor (Invitrogen). After phenol/chloroform extraction, Terminator-digested RNA was precipitated with ethanol, and then resuspended in nuclease-free H_2_O.

Libraries were prepared from the rRNA-depleted or Terminator-digested RNAs using the TruSeq Stranded mRNA Sample Preparation Kit (Illumina) and the IDT for Illumina – TruSeq RNA UD indexes (Illumina). Paired-end sequencing (2 x 50 nt) was performed on a NovaSeq 6000 system (Illumina).

### Total-Seq data processing and analysis

Reads were trimmed using Trim Galore (https://github.com/FelixKrueger/TrimGalore) and mapped on the S288C reference genome (R64-2-1, including the 2-micron plasmid), with addition of either ERCC RNA spike-in sequences or the *Schizosaccharomyces pombe* genome (ASM294v2, for the heat-shock dataset) using version 2.2.0 of Hisat (43), with default parameters and a maximum size for introns of 5000. All subsequent analyses used uniquely mapped reads. Mapping statistics are presented in Table S5.

Gene counts were computed using version 2.0.0 of featureCounts (44), and then normalized using the estimateSizefactorsForMatrix function from the DESeq2 package (45). Tag densities were obtained as: normalized gene count/gene length.

For all the RNA-Seq data produced in this study, normalization on the ERCC RNA spike-in signal was used in a first time to control that snoRNAs expression is not affected in the mutant/condition analysed, and snoRNA counts were then used for normalization, as previously described (39,46).

For the heat-shock dataset (retrieved from the NCBI Gene Expression Omnibus using accession number GSE148166), gene counts were normalized on the *S. pombe* spike-in RNA, as snoRNAs levels were abnormally low in both stressed and control cells, probably due to differences in the library preparation protocol (47).

XUTs were defined as up-regulated in a given condition when showing a >2-fold enrichment in this condition *vs* the control, with a *P*-value <0.05 (adjusted for multiple testing with the Benjamini-Hochberg procedure) upon differential expression analyses using DESeq2 (45).

### Ribo-Seq libraries preparation

Ribo-Seq analysis was performed from two biological replicates YAM1 (WT) and YAM202 (*upf1Δ*) cells, grown to mid-log phase (OD_600_ 0,5) at 30°C in YPD, then treated or not for 15 minutes with CHX (100 µg/ml, final concentration). For each sample, 250 ml of cells were harvested by centrifugation at room temperature and directly frozen in liquid nitrogen after supernatant removal.

Cells were lysed in 1X lysis buffer (10 mM Tris-HCl pH 7.4, 100 mM NaCl, 30 mM MgCl_2_) supplemented by 2X cOmplete Protease Inhibitor Cocktail (Roche) and ribosome protected fragments (RPFs) were prepared as previously described (48), except that the purification on sucrose cushion was performed before the digestion with RNase I (Ambion, 5 units/UA_260_). Biotinylated oligonucleotides (IDT - Integrated DNA Technologies) used for ribo-depletion are listed in Table S4.

Libraries were then prepared from 10 ng of RPFs using the D-Plex Small RNA-Seq kit for Illumina (Diagenode) and the D-Plex Unique Dual Indexes for Illumina – set A (Diagenode). The RPFs were diluted in a final volume of 8 µl before the addition of 2 µl of Dephoshorylation Buffer, 5 µl of Crowding Buffer and 0.5 µl of Dephosphorylation Reagent. The samples were incubated for 15 minutes at 37°C. RNA tailing was performed by adding 1.5 µl of Small Tailing Master Mix (1 µl of Small Tailing Buffer + 0.5 µl of Small Tailing Reagent) to the dephosphorylated RNAs, and incubating the samples for 40 minutes at 37°C. The samples were transferred on ice for 2 minutes before the addition of 1 µl of the Reverse Transcription Primer (RTPH). The samples were denaturated for 10 minutes at 70°C and then cooled down to 25°C at a 0.5°C/sec rate. A Reverse Transcription Master Mix (RTMM) was prepared by mixing 5 µl of Reverse Transcription Buffer and 1 µl of Reverse Transcription Reagent; 6 µl of this mix were added to the samples, which were then incubated for 15 minutes at 25°C. After adding 2 µl of Small Template Switch Oligo, the samples were incubated for 120 minutes at 42°C, then heated for 10 minutes at 70°C and finally kept at 4°C. For the PCR amplification, 20 µl of D-Plex Primer UDI and 50 µl of PCR Master Mix were added, then the following program was run: initial denaturation at 98°C for 30 seconds; 10 cycles including 15 seconds at 98°C followed by 1 minute at 72°C; final incubation of 10 minutes at 72°C; hold at 4°C. The libraries were then purified using the Monarch PCR & DNA Cleanup Kit (NEB), using a 5:1 ratio of Binding Buffer: Sample. Purified DNA was eluted in 50 µl of nuclease-free H_2_O (Ambion). A second cleanup of the libraries was performed using 1 volume of AMPure XB beads (Beckman). Libraries were eluted in 20 µl of nuclease-free H_2_O (Ambion), and then quantified using the Qubit dsDNA HS assay (Invitrogen). Finally, the size and the molarity of each library were determined using a High Sensitivity D1000 ScreenTape in a 4200 TapeStation (Agilent Technologies).

Single-end sequencing (50 nt) of the libraries was performed on a NovaSeq 6000 system (Illumina).

### Detection of translated XUTs/smORFs using Ribotricer

Unique molecular identifiers (UMI) were extracted using umi_tools (49), and then used to discard PCR duplicates. Reads were trimmed using cutadapt v2.10 (50), and then mapped using Hisat v2.0.0 (43), as above. Reads mapping on rRNA were discarded. Subsequent analyses only used uniquely mapped reads with a size comprised between 25 and 35 nt. Mapping statistics are presented in Table S5.

Ribotricer 1.3.1 was used to extract translated ORFs (minimum length of 15 nt) based on *S. cerevisiae* genome annotation (including XUTs), using ATG as the start codon and a phase-score cutoff of 0.318, as recommended by the authors (51). The phasing of Ribo-Seq data was also analysed independently (see Figure S8). List 1 of translated XUTs was obtained after pooling the bam files from all conditions. List 2 was obtained by analyzing each condition separately, pooling the bam files from the two biological replicates. List 3 was obtained from list 2, upon application of a coverage filter (at least 10 reads per translated smORF).

### Protein extraction and Western blot

Protein extracts were prepared from exponentially growing cells, using a standard method based on cell lysis with glass beads in ‘IP’ buffer (20 mM HEPES pH 7.5, 100 mM NaCl, 0.5 mM EDTA, 1 mM DTT, 20% glycerol), supplemented with 0.05% NP40, 0.5X cOmplete Protease Inhibitor Cocktail (Roche) and 1 mM AEBSF.

40 µg of total extracts were separated on NuPAGE 4-12% Bis-Tris gel (Invitrogen) in 1X NuPAGE MOPS SDS running buffer (Invitrogen), and then transferred on a nitrocellulose membrane using iBlot 2 Transfer Stack system (Invitrogen), with program ‘0’.

The FLAG-tagged peptide and Pgk1 were detected using the anti-FLAG M2 (Sigma; cat# F1365; 1:1000 dilution) and anti-Pgk1 22C5D8 (abcam; cat# ab113687; 1:10000 dilution) monoclonal primary antibodies, revealed using the SuperSignal West Femto Maximum Sensitivity Substrate (Thermo Scientific) and the SuperSignal West Pico Chemiluminescent Substrate (Thermo Scientific), respectively, with a ChemiDoc Imaging System (BioRad). The secondary antibody was the anti-mouse IgG (whole molecule)–peroxidase antibody produced in rabbit (Sigma; cat# A9044; 1:10000 dilution).

### Mass spectrometry (MS)

MS was performed starting from crude extracts of *upf1Δ* (YAM202) cells grown to mid-log phase at 30°C in in CSM medium with 0.1% proline as nitrogen source and containing 0.003% SDS, and then treated for 3 hours with 50 µM MG-132 proteasome inhibitor (Sigma: cat# M7449).

The crude extracts were separated on 4-12% bis-tris gels (Invitrogen; cat# NP0326BOX) in MES buffer (Invitrogen; cat# NP0002), then the region of the gel encompassing the small peptides fraction (1-10 kDa) was cut in bands of about 2 mm and subjected to in-gel trypsin digestion as previously described (52) before submission to MS analysis. Trypsin-generated peptides were analyzed by nanoLC–MSMS using a nanoElute liquid chromatography system (Bruker) coupled to a timsTOF Pro mass spectrometer (Bruker). Peptides were loaded on an Aurora analytical column (ION OPTIK, 25 cm × 75 m, C18, 1.6 m) and separated with a gradient of 0–35% of solvent B for 100 minutes. Solvent A was 0.1% formic acid and 2% acetonitrile in water and solvent B was acetonitrile with 0.1% formic acid. MS and MS/MS spectra were recorded from m/z 100 to 1700 with a mobility scan range from 0.6 to 1.4 V s/cm2. MS/MS spectra were acquired with the PASEF (Parallel Accumulation Serial Fragmentation) ion mobility-based acquisition mode using a number of PASEF MS/MS scans set as 10. MS and MSMS raw data were processed and converted into mgf files with DataAnalysis software (Bruker). Protein identifications were performed using the MASCOT search engine (Matrix Science, London, UK) against SwissProt and a non-canonical proteins homemade database. Database searches were performed using trypsin cleavage specificity with two possible missed cleavages. Carbamidomethylation of cysteines was set as fixed modification and oxidation of methionines as variable modification. Peptide and fragment tolerances were set at 10 ppm and 0.05 Da, respectively. Only ions with a score higher than the identity threshold and a false-positive discovery rate of less than 1% (Mascot decoy option) were considered.

As a control, we included a synthetic, labelled AQUA version of the peptide derived from *XUT0741* (AQUA Basic Heavy grade, Thermo Scientific). The sequence of the synthetic peptide was MPY(I)TNTAEATMSTV, where (I) corresponds to a stable isotope isoleucine (+7Da).

## RESULTS

### Translation determines the decay of cytoplasmic NMD-sensitive lncRNAs

The NMD-sensitivity of XUTs suggests that translation determines their decay. We anticipated that inhibiting translation would result in the accumulation of NMD-sensitive XUTs. To explore this idea, we treated exponentially growing WT cells with cycloheximide (CHX), a translation elongation inhibitor which binds the E site of the ribosome and prevents tRNA release and ribosome translocation (53). Samples were collected at different time points after addition of the drug, then total RNA was extracted and analysed by Northern blot. We observed that *XUT1678* and *XUT0741*, two NMD-sensitive XUTs that we previously characterized (39), accumulate as soon as 5-10 min after CHX addition (Figure 1A). This effect is reversible, as the levels of both lncRNAs decreased after washing the CHX-treated cells and returning them to growth in fresh medium without CHX (Figure 1B). We noted that the 5’ ITS1 fragment, a well-known physiological target of Xrn1 (54), did not accumulate in CHX-treated WT cells (Figure 1A), indicating that CHX does not block the activity of Xrn1. In addition, we found that anisomycin (ANS), which also inhibits translation elongation but at a different stage than CHX (Figure S1A), led to a similar accumulation of *XUT1678* and *XUT0741* in WT cells (Figure 1C), reinforcing our hypothesis of a general translation-dependent lncRNAs decay process.

**Figure 1.**
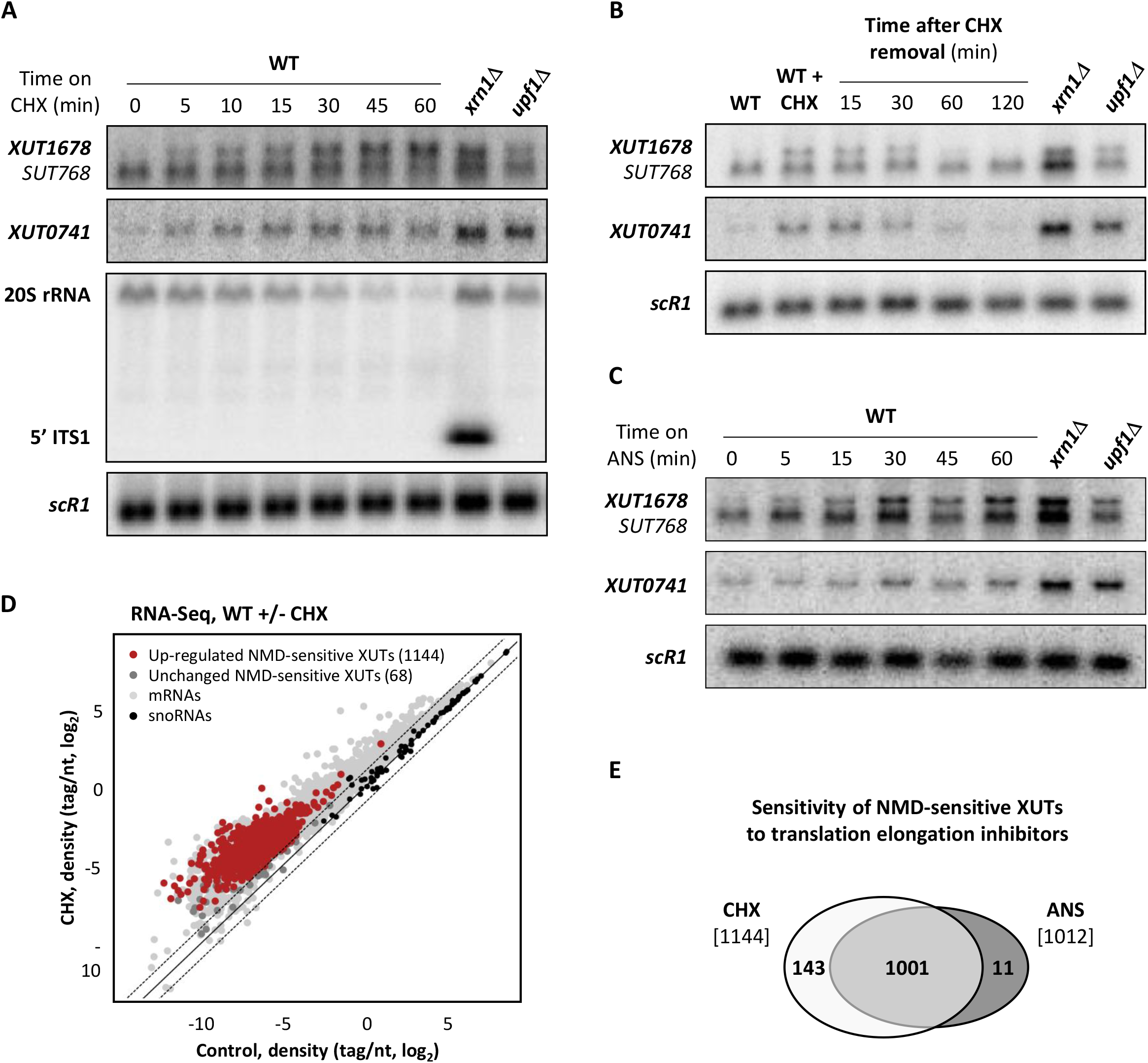
NMD-sensitive lncRNAs accumulate upon translation inhibition. **A.** WT (YAM1) cells were grown to mid-log phase in rich (YPD) medium at 30°C. CHX was then added at a final concentration of 100 μg/ml, and samples were collected at different time points. Untreated *xrn1Δ* (YAM6) and *upf1Δ* (YAM202) cells, grown under the same conditions, were used as controls. Total RNA was extracted and analysed by Northern blot. *XUT1678* (and the overlapping *SUT768*), *XUT0741*, the 5’ ITS1 fragment (as well as the 20S pre-rRNA it derives from) and *scR1* (loading control) were detected using ^32^P-labelled AMO1595, AMO1762, AMO496 and AMO1482 oligonucleotides, respectively. **B.** WT (YAM1), *xrn1Δ* (YAM6) and *upf1Δ* (YAM202) cells were grown as above. CHX was then added to the WT cells for 15 min (100 μg/ml, final concentration). The CHX-treated cells were then washed with fresh pre-heated YPD medium and re-incubated at 30°C. Samples of washed cells were collected after 15, 30, 60 and 120 min. Total RNA was extracted and analysed by Northern blot as described above. **C.** Same as Figure 1A using ANS (100 μg/ml final concentration) instead of CHX. **D.** Total RNA-Seq was performed using total RNA extracted from exponentially growing WT (YAM1) cells (grown as above) treated for 15 min with CHX (100 μg/ml, final concentration) or with an equal volume of DMSO (control). The scatter plot shows the RNA-Seq signals (tag densities, log_2_ scale) for the NMD-sensitive XUTs, mRNAs (light grey dots) and snoRNAs (black dots) in CHX-treated and control WT cells. The significantly up-regulated (CHX/control fold-change >2, *P*-value <0.05) and unaffected NMD-sensitive XUTs are represented as red and dark grey dots, respectively. **E.** Venn diagram showing the number of NMD-sensitive XUTs that accumulate in CHX- and/or ANS-treated WT cells.

These results were extended at the genome-wide level using RNA-Seq, showing that the majority of NMD-sensitive XUTs significantly accumulate (fold-change >2, *P*-value <0.05) in WT cells treated with CHX or ANS (Figure 1D-E; see also Figure S1B and Table S1). In contrast, CHX and ANS only had a moderate effect on Cryptic Unstable Transcripts (CUTs), which are degraded in the nucleus by the Exosome (55–57), indicating that translation primarily impacts cytoplasmic transcripts (Figure S1C).

The observation that NMD-sensitive XUTs rapidly accumulate in WT cells following inhibition of translation elongation is consistent with the idea that translation determines the degradation of cytoplasmic NMD-sensitive lncRNAs.

### Translation also affects lncRNAs decay independently of NMD

While NMD targets most XUTs, about 30% of them remain NMD-insensitive (39). We asked whether these cytoplasmic transcripts that escape NMD also react to translation elongation inhibition. Our RNA-Seq data showed that most NMD-insensitive XUTs significantly accumulate in CHX- and ANS-treated WT cells (Figure 2A-B, see also Figure S2A and Table S1). This indicates that translation can affect decay of XUTs independently of NMD.

**Figure 2.**
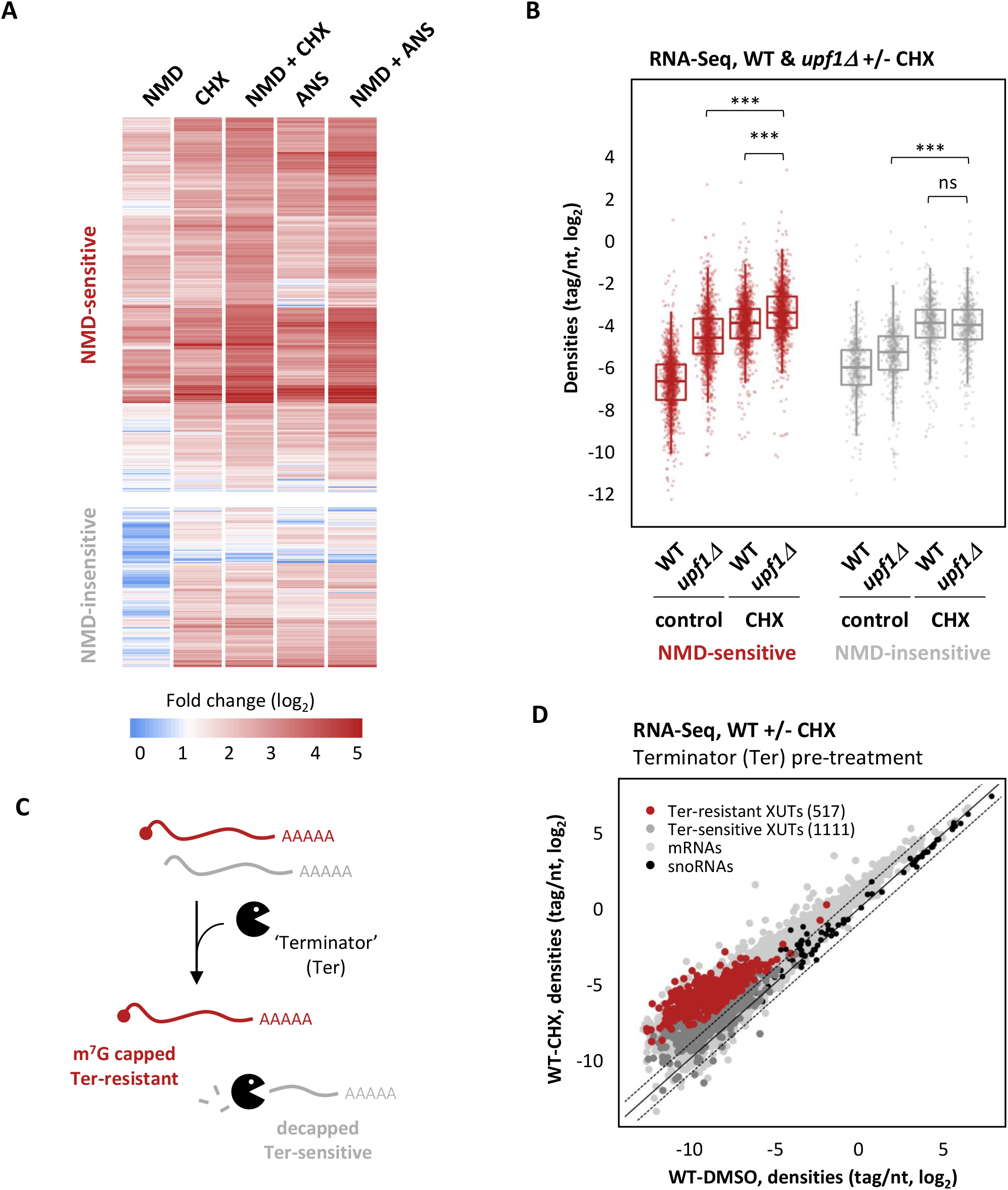
Translation also impacts XUTs independently of NMD. **A.** Total RNA-Seq was performed in WT (YAM1) and *upf1Δ* (YAM202) cells, with or without treatment with CHX (15 min, 100 μg/ml final concentration) or ANS (30 min, 100 μg/ml final concentration). Densities were computed for NMD-sensitive and NMD-insensitive XUTs, using our previously published annotation (39). The sensitivity to NMD and/or CHX/ANS of each transcript is shown as an heatmap of the fold-change (log_2_ scale) relative to the corresponding control WT cells (treated for the same time with an equal volume of DMSO). Note in the first column that some XUTs previously annotated as NMD-sensitive were not detected as such in this novel dataset (see Table S1). **B.** Same as above. The data are presented as densities (tag/nt, log_2_ scale) for NMD-sensitive and NMD-insensitive XUTs in control (DMSO) or CHX-treated WT (YAM1) and *upf1Δ* (YAM202) cells. *** *P*-value < 0.001; ns, not significant upon two-sided Wilcoxon rank-sum test (adjusted for multiple testing with the Benjamini–Hochberg procedure). **C.** Schematic representation of the action of the Terminator 5’-phosphate-dependent exonuclease, which degrades RNAs that are decapped (grey), but not those with an intact m^7^G cap (red). **D.** Total RNA-Seq was performed using the same RNA extracts as in Figure 1D, including a treatment with the Terminator 5’-phosphate-dependent exonuclease before the preparation of the libraries. The data are presented as in Figure 1D, the red dots representing the 517 CHX-sensitive XUTs that are still detected as significantly up-regulated in CHX-treated WT cells (CHX/control fold-change >2, *P*-value <0.05) upon Terminator treatment. The other XUTs (Terminator-sensitive) are represented as dark grey dots.

To further explore this idea, we performed RNA-Seq in CHX-treated *upf1Δ* cells. This revealed that NMD inactivation and CHX have a synergistic effect on NMD-sensitive XUTs (Figure 2B), their global levels being significantly higher in the CHX-treated *upf1Δ* cells compared to the untreated *upf1Δ* cells (*P* < 2^e-26^, Wilcoxon rank-sum test) or the CHX-treated WT cells (*P* < 2^e-26^, Wilcoxon rank-sum test). Similar observations were made with ANS (see Figure S2A). Importantly, this synergy between NMD inactivation and CHX- or ANS-induced translation elongation inhibition was only observed for the NMD-sensitive XUTs, but not for the NMD-insensitive ones (Figure 2B, see also Figure S2A).

These observations raise the question of the mechanism by which translation affects XUTs independently of NMD. In a previous study, CHX has been proposed to interfere with the decapping of the *MFA2* mRNA, leading to its stabilization (41). This led us to assess whether this could also be the case for the CHX-sensitive XUTs.

To determine the capping status of the XUTs that accumulate upon CHX treatment, we performed RNA-Seq using the same RNA extracts as above, but included a treatment with the Terminator 5’-phosphate-dependent exonuclease, which degrades RNAs with 5’-monophosphate ends but not those with an intact m^7^G cap (Figure 2C). This revealed that 517 (35%) of the XUTs that accumulate in CHX-treated WT cells are Terminator-resistant, indicating that they accumulate as capped RNAs (Figure 2D; see also Table S1).

Decapping and NMD are functionally linked (58,59). According to the current models, the recruitment of the NMD core factors precedes the recruitment of the decapping machinery (60). As NMD depends on translation, one could imagine that NMD is less efficient in CHX-treated cells, which would in turn negatively impact the recruitment of the decapping factors. To explore how NMD inactivation affects the decapping of XUTs, we assessed the Terminator-sensitivity of XUTs in *upf1Δ* cells. Unexpectedly, we found that most NMD-sensitive XUTs accumulating in the *upf1Δ* mutant are decapped, as shown by their global sensitivity to the Terminator exonuclease (Figure S2B-C). In fact, only 149 XUTs significantly accumulate in the *upf1* mutant (*upf1Δ*/WT ratio >2, *P*-value < 0.05) following Terminator digestion (Figure S2B; see also Table S1).

Thus, since most XUTs are efficiently decapped in the absence of Upf1, their accumulation in the NMD mutant is unlikely to reflect a decapping defect, but rather the inability of Xrn1 to access them. In addition, since the effect of CHX on the decapping of XUTs is more important than the effect of NMD inactivation, we conclude that the physical presence of elongating ribosomes can interfere with the decapping of a fraction of XUTs, independently of NMD.

### LncRNAs levels remain globally unchanged upon stress-induced inhibition of translation initiation

The data described above show that treating WT cells with CHX or ANS results in the accumulation of most XUTs. At the molecular level, these drugs act by arresting elongating ribosomes on their RNA substrates, a property which is widely exploited in Ribo-Seq analyses (61,62).

Interestingly, mRNA degradation is known to occur co-translationally (63), and several reports have proposed that the physical presence of ribosomes on an mRNA can interfere with its co-translational degradation by Xrn1 (64,65). This led us to investigate whether the accumulation of XUTs observed in the presence of CHX or ANS could reflect a protective effect of the ribosomes themselves, forming a physical obstacle that would block Xrn1 (Figure 3A). According to this model, XUTs should not accumulate when ribosomes are not pre-loaded on the transcripts, *i.e.* in conditions where translation initiation is inhibited (Figure 3A).

**Figure 3.**
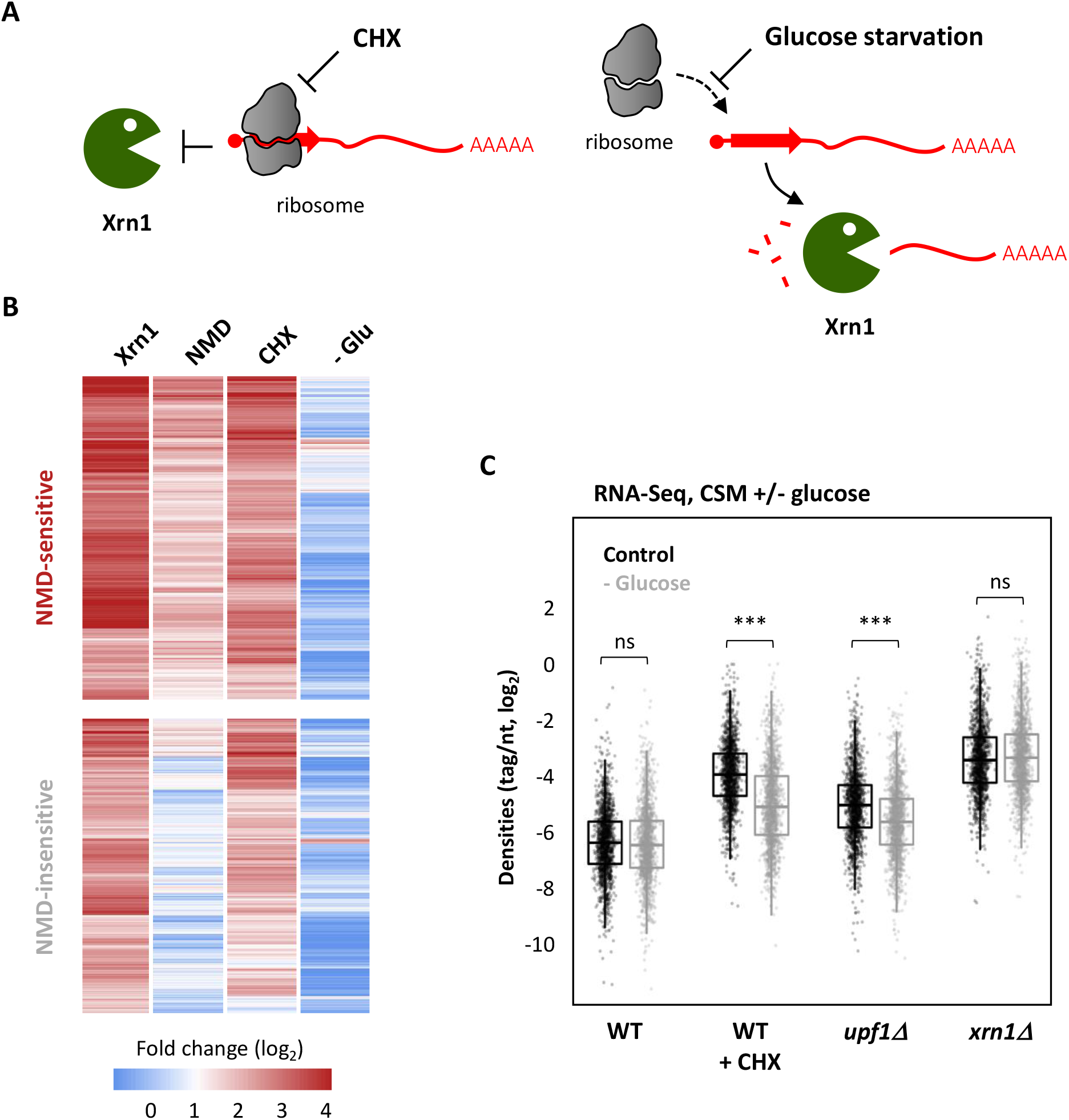
XUTs levels remain unaffected upon translation initiation inhibition. **A.** Working model showing the presumed effect of CHX-mediated inhibition of translation elongation (left) and of stress-induced inhibition of translation initiation (glucose starvation, right) on Xrn1-dependent degradation of XUTs (red). The red arrow on the XUT represents a smORF. **B.** Total RNA-Seq was performed in WT (YAM1), *xrn1Δ* (YAM6) and *upf1Δ* (YAM202) grown in CSM. WT cells grown in the same conditions and then submitted to a CHX treatment or glucose starvation (-Glu) were also included. Densities (tag/nt) were computed for the 1470 XUTs significantly up-regulated in the *xrn1* mutant grown in CSM (see Figure S3A), which were then separated according to their sensitivity to NMD (see Figure S3B). The sensitivity of each of these XUTs to CHX and glucose starvation is presented as an heatmap of the fold-change (log_2_ scale). As an indication, the sensitivity of these XUT to Xrn1 (*xrn1Δ*/WT) and NMD (*upf1Δ*/WT) is also presented. **C.** Box-plot showing the RNA-Seq signals (densities, tag/nt, log_2_ scale) for the same set of XUTs as in panel B, in WT (YAM1), *upf1Δ* (YAM202) and *xrn1Δ* (YAM6) cells grown in CSM with glucose (control; black) or undergoing glucose starvation for 16 min (-Glucose; grey). An aliquot of WT cells was then treated with CHX for 15 min. *** *P*-value < 0.001; ns, not significant upon two-sided Wilcoxon rank-sum test (adjusted for multiple testing with the Benjamini–Hochberg procedure).

Glucose is the preferred source of carbon of yeast. Different works have shown that glucose depletion results in a global inhibition of translation at the initiation level, with a rapid loss of polysomes (66–68). More recently, the Tollervey lab showed that the stress response induced upon glucose starvation or heat-shock is associated with a displacement of translation initiation factors from mRNAs (47).

To study the impact of translation initiation inhibition on XUTs, we performed RNA-Seq using WT cells grown in glucose-containing medium and then shifted for 16 min into medium containing glycerol and ethanol (see Figure S3A-C). Strikingly, XUTs levels remain globally unchanged upon glucose depletion, which is in sharp contrast with the effect of translation elongation inhibition (CHX) observed in normal (glucose-containing) medium (Figure 3B; see also Table S1). Similarly, a re-analysis of RNA-Seq data obtained in heat-shock conditions (47) showed that this stress does not lead to a global accumulation of XUTs either (see Figure S3D and Table S1).

Importantly, we observed that the sensitivity of XUTs to CHX and to NMD significantly decreases upon glucose depletion (Figure 3C), confirming that translation is inhibited in this condition. In contrast, the sensitivity of XUTs to Xrn1 was unchanged (Figure 3C), indicating that Xrn1-dependent degradation of XUTs remains fully functional upon glucose depletion.

Altogether, these data suggest that the stabilization of XUTs observed in CHX-treated WT cells is mediated by the elongating ribosomes, sterically protecting them from degradation. In contrast, when they are not bound by ribosomes, XUTs are by default degraded by Xrn1.

### Translational Landscape of yeast lncRNAs

A previous Ribo-Seq analysis in *upf1Δ* yeast cells revealed 47 smORFs on 43 lncRNAs, providing a first proof-of-principle that lncRNAs can also be bound by ribosomes in *S. cerevisiae* (15). In order to define a more comprehensive translational landscape of yeast lncRNAs, we performed a new Ribo-Seq experiment in WT and *upf1Δ* cells, producing two datasets for each genetic background: one in native conditions (untreated cells), and a second using CHX-treated cells (Figure 4A).

**Figure 4.**
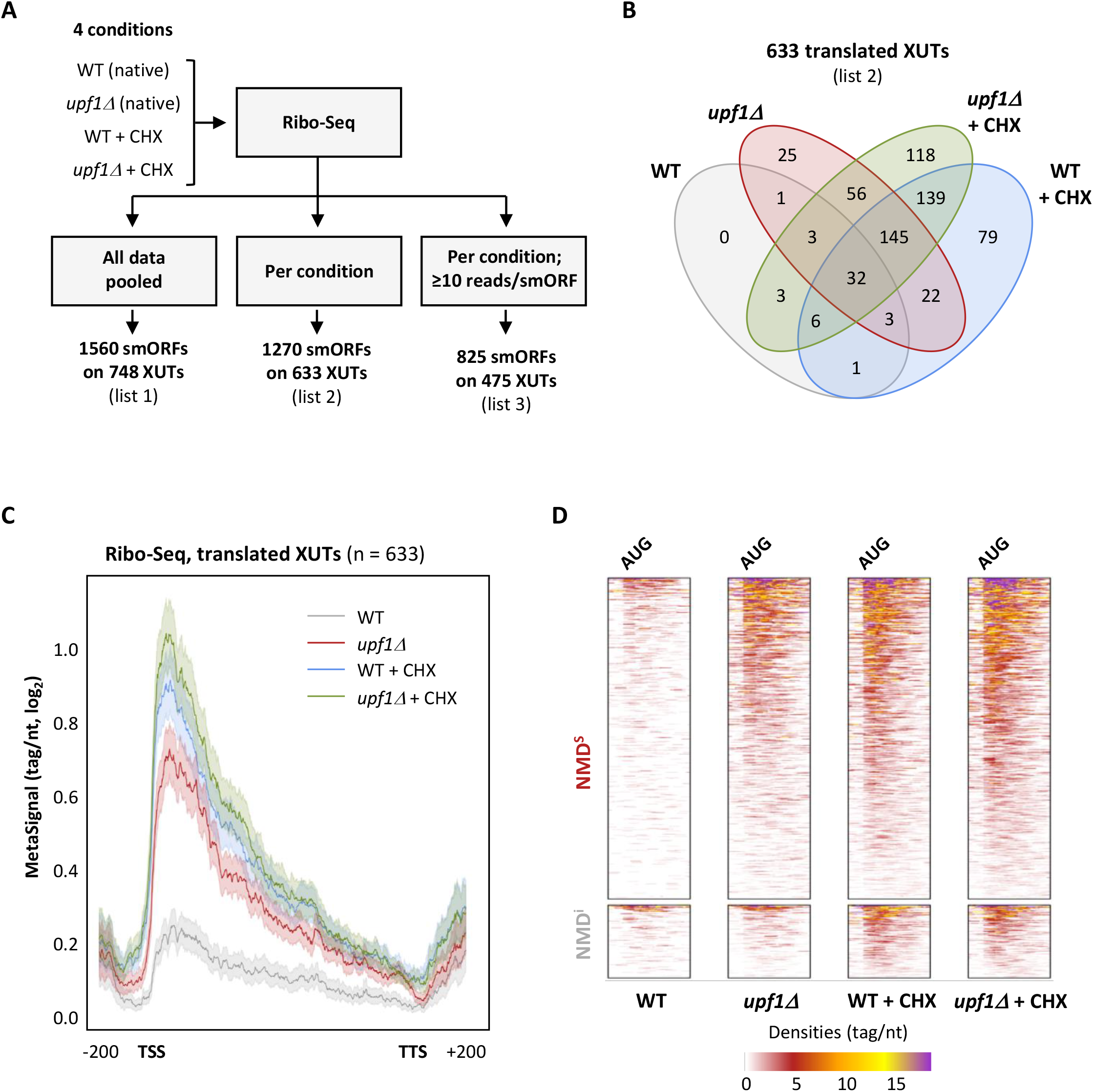
Translational landscape of XUTs. **A.** Experimental scheme. Ribo-Seq libraries were prepared from biological duplicates of WT and *upf1Δ* cells grown in native conditions or treated for 15 min with CHX (100 μg/ml final concentration). SmORFs (≥ 5 codons, starting with an AUG) were detected using the ribotricer software (51), pooling all conditions together (list 1) or analyzing them separately (list 2). A third list was produced from list 2 upon application of a signal threshold (≥ 10 reads/smORF). See lists in Table S2. **B.** Venn diagram showing the number of XUTs detected as translated by Ribotricer (list 2) in each of the indicated conditions. See also Table S2. **C.** Metagene of Ribo-Seq signals along the 633 translated XUTs (list 2). For each condition, the densities (tag/nt, log_2_) along the XUTs +/- 200 nt were piled up, then the average signal was plotted. The shading surrounding each line denotes the 95% confidence interval. **D.** Heatmap view of the Ribo-Seq signals (densities, tag/nt) from positions −50 to +150 relative to the AUG codon of the smORF showing the highest signal for the 510 NMD-sensitive and 123 NMD-insensitive XUTs detected as translated. A separate heatmap is shown for each condition.

As a first approach, we pooled our Ribo-Seq data and searched for smORFs (≥ 5 codons, starting with an AUG codon) using Ribotricer (51), which directly assesses the 3-nt periodicity of Ribo-Seq data to identify actively translated ORFs (see Materials & Methods). This led to the identification of 1560 translated smORFs on 748 XUTs (Figure 4A; list 1 in Table S2). We then repeated the same procedure, separating the conditions, which produced a refined list of 1270 smORFs from 633 XUTs, translated in at least one condition (Figure 4A-B; list 2 in Table S2). Applying an additional coverage threshold (≥ 10 reads/smORF in at least one condition) restricted the list to 825 smORFs for 475 XUTs (Figure 4A; list 3 in Table S2), which corresponds to the most robust candidates within the set of translated smORFs/XUTs, showing the highest levels of translation and being translated in at least one condition. However, since the translation of lncRNAs could also be transient and occur at low levels, we decided to use the second list of 633 translated XUTs as a compromise for the descriptive analysis below. Figure 4C shows a metagene view of the Ribo-Seq signals for these XUTs in the four conditions. A similar metagene analysis for the other XUTs (not detected as translated) revealed that the signals are globally lower, suggesting that our analysis captured the XUTs displaying the highest levels of translation (Figure S4A).

Of these 633 XUTs, 510 are NMD-sensitive and 123 are NMD-insensitive (Figure 4D; see also Table S2). Notably, 297 of them are detected as translated in the native condition, essentially in the *upf1Δ* mutant (Figure 4B). As one would expect, combining NMD inactivation and CHX treatment strongly improves the detection of XUTs translation (Figure 4B). Cumulatively, 411 XUTs were detected as translated in at least two datasets (Figure 4B).

The smORFs detected on XUTs display a median size of 87 nt (Figure S4B), which is in line with the size of non-canonical (n)ORFs recently identified in yeast and the fact that these nORFs are globally shorter than canonical ORFs (69).

We noted that for half of the XUTs (311/633), Ribotricer detected more than one smORF (Figure S4C). This could reflect the potential of several smORFs on a same XUT to attract the translation machinery and/or the existence of distinct isoforms for a same XUT, possibly displaying different boundaries and encompassing different smORFs. Interestingly, for 75% of the translated XUTs, the smORF showing the highest Ribo-Seq signal corresponds to one of the first three smORFs predicted in the XUTs sequence (Figure S4D). This is consistent with the observation that ribosomes preferentially bind the 5’-proximal region of the translated XUTs (Figure 4C). Note that this profile is unlikely to be an artifact due to the CHX treatment, as it is also observed in native conditions (Figure 4C).

Together, these data show that a substantial fraction of XUTs carry smORFs that are experimentally detected as actively translated.

### Features of translated lncRNAs

The identification of translated smORFs on XUTs prompted us to investigate the potential regulatory effect on XUTs decay.

As described above, 517 XUTs accumulate as capped RNAs (Terminator-resistant) in CXH-treated cells (Figure 2D). Notably, 300 of them (58%) are detected as translated in our Ribo-Seq analysis, which is significantly more than expected by chance (*P* = 3.26 e-29, Chi-square test of independence; Figure S5A-B). XUTs translation therefore correlates with their accumulation as capped transcripts when the elongating ribosomes are blocked on their template.

We speculated that ribosomes could sterically block Dcp2 because the translated smORF of these XUTs is close to the transcript start site (TSS). To investigate this possibility, we computed the 5’ UTR length for each translated XUTs. Surprisingly, we found no significant difference between the Terminator-resistant and the Terminator-sensitive XUTs (Figure S5C). Furthermore, ribosome occupancy along the 5’ region was similar between the two subgroups of XUTs (Figure 5A; see also Figure S5D). Thus, the accumulation of capped XUTs upon translation elongation inhibition is not the consequence of a higher ribosome density along the TSS-proximal region.

**Figure 5.**
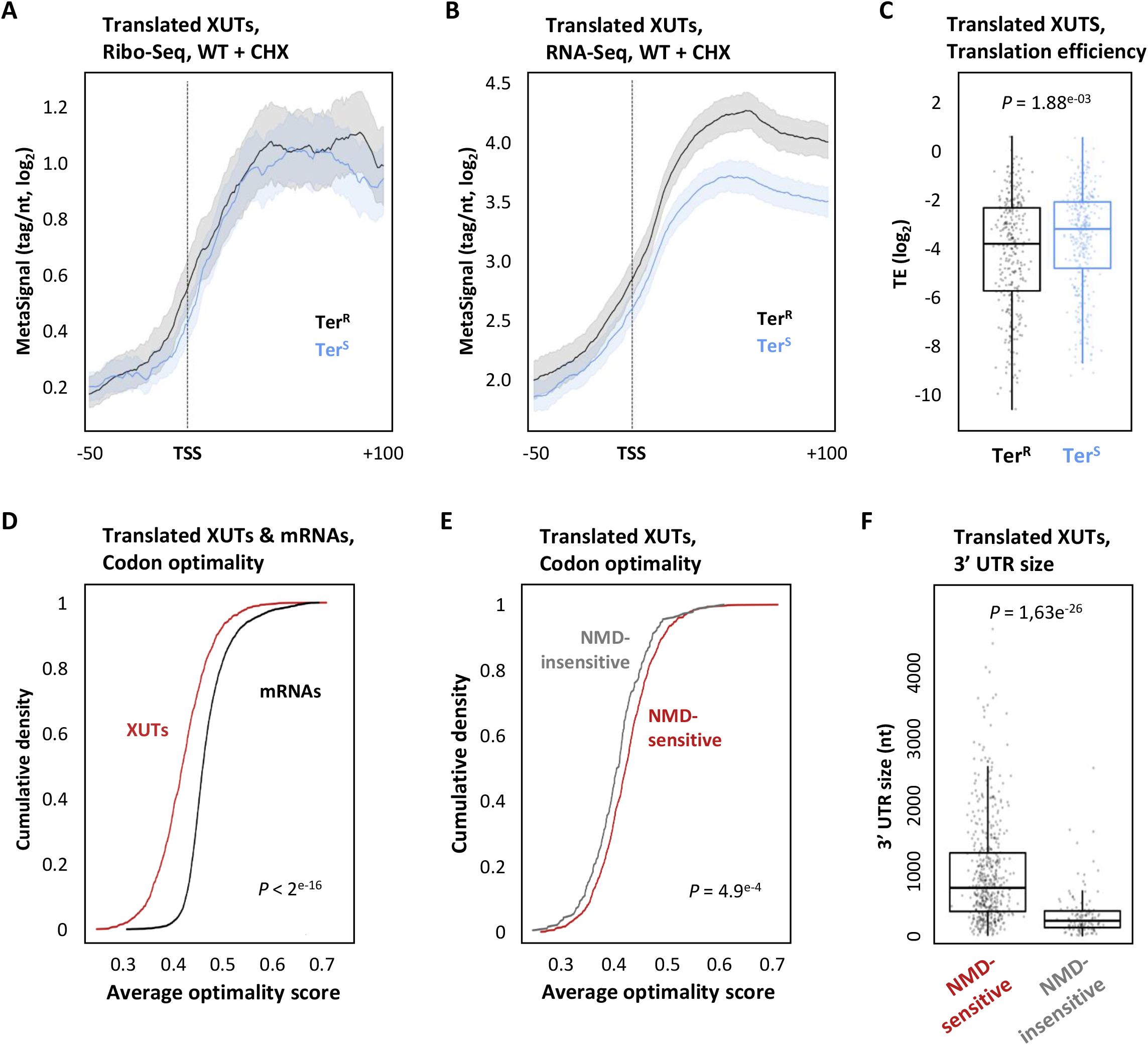
Features of translated XUTs. **A.** Metagene of Ribo-Seq signals along positions −50 to +100 relative to the TSS of the 300 Terminator-resistant (Ter^R^, black) and the 333 Terminator-sensitive (Ter^S^, blue) translated XUTs. The metagene was produced as described in Figure 4C, using signals obtained in CHX-treated WT cells. **B.** Same as above, using RNA-Seq signals obtained in the same condition. **C.** Box-plot showing the translation efficiency (TE) for the Terminator-resistant and Terminator-sensitive translated XUTs. TE for each XUT was computed by normalizing the Ribo-Seq signals along the TSS to TSS+50 region in CHX-treated cells to the RNA-Seq signals in the same region and conditon. The *P*-value obtained upon two-sided Wilcoxon rank-sum test is indicated. **D.** Average codon optimality score for the 633 translated XUTs (red) *vs* mRNAs (noir), shown as a cumulative frequency plot. For XUTs with several annotated smORFs, we considered the smORF displaying the highest Ribo-Seq signal. The indicated *P*-value was obtained upon Kolmogorov–Smirnov test. **E.** Same as above for the 510 NMD-sensitive (red) and 123 NMD-insensitive (grey) translated XUTs. **F.** Box-plot showing the size of the 3’ UTR for the NMD-sensitive and NMD-insensitive translated XUTs. As in panel D, when several translated smORFs have been identified for a same XUT, we considered the smORF with the highest Ribo-Seq signal to compute the 3’ UTR length. The *P*-value was obtained upon two-sided Wilcoxon rank-sum test.

The translated XUTs are globally more abundant than the other XUTs in CHX-WT cells (Figure S5E), and we observed a positive correlation between XUTs translation and abundance, which is even more pronounced for the Terminator-sensitive XUTs (Figure S5F). Yet, the Terminator-resistant XUTs are globally more abundant than the Terminator-sensitive ones (Figure 5B; see also Figure S5G), as one could expect since the latter are decapped and can therefore be directly degraded by Xrn1. Since ribosome density was similar between the two subgroups of XUTs, the Terminator-sensitive XUTs display a higher translation efficiency (Figure 5C; *P* = 1.88^e-03^, Wilcoxon rank-sum test), computed by normalizing the Ribo-Seq signal to the transcript abundance. In other words, this result indicates that translating a XUT with a lower efficiency negatively impacts its decapping.

Then, we analysed the codon optimality for the translated XUTs. Codon optimality affect translation elongations and has been associated to mRNA stability (70,71). In fact, unstable mRNAs are enriched in non-optimal codons, which are supposed to be decoded less efficiently (70). In addition, ‘normal’ mRNAs (*i.e*. devoid of premature termination codon) yet targeted by NMD are also enriched in non-optimal codons (34). This prompted us to determine the codon optimality score of the translated XUTs. Globally, we observed that it is significantly lower than for mRNAs (Figure 5D; *P* < 2^e-16^, Kolmogorov–Smirnov test). However, while NMD sensitivity correlates with a lower codon optimality in the case of mRNAs (Figure S5H; *P* = 1.1^e-16^, Kolmogorov–Smirnov test), XUTs display an opposite pattern, the average codon optimality score being significantly higher for the NMD-sensitive XUTs (Figure 5E; *P* = 1.1^e-16^, Kolmogorov–Smirnov test).

Finally, we computed the size of the 3’ UTR of the translated XUTs and observed that it is significantly longer for the NMD-sensitive XUTs than for the NMD-insensitive XUTs (Figure 5F; median = 733 nt *vs* 236 nt; *P* = 1.63e^-26^, Wilcoxon rank-sum test), strongly indicating that the length of the 3’ UTR is a key determinant for XUTs degradation by NMD, as for mRNAs (33,34).

In summary, these results highlight a correlation between the translation efficiency of XUTs and their decapping. They also reveal that the NMD-sensitive XUT display a higher codon optimality score and a longer 3’ UTR in comparison to the NMD-insensitive ones.

### The long 3’ UTR of *XUT0741* is a major determinant of its NMD-sensitivity

For yeast mRNAs, the length of the 3’ UTR is known to be critical for NMD activation (33–35). The observation that the 3’ UTR is significantly longer for the NMD-sensitive XUTs compared to NMD-insensitive XUTs suggests that it might also constitute a key determinant of the NMD-sensitivity for XUTs, as for mRNAs (33,34). We therefore investigated this hypothesis, using the NMD-sensitive *XUT0741* as a model candidate.

*XUT0741* belongs to the top list of translated XUTs, with a single 5’-proximal smORF (15 codons), detected by each of our different analyses (see Figure S6A and Table S2). This smORF is followed by a 1.3 kb long 3’ UTR containing multiple stop codons in the same frame (Figure 6A). To explore the role of the 3’ UTR as a *cis* element determining its NMD-sensitivity, we designed six mutants of *XUT0741* by mutating several of these in-frame stop codons, thereby progressively lengthening the smORF while shortening the 3’ UTR (Figure 6A; see sequences in Supplementary File 1). These mutant alleles were integrated at the genomic locus in WT and *upf1Δ* strains, then their expression and NMD-sensitivity were assessed by strand-specific RT-qPCR.

**Figure 6.**
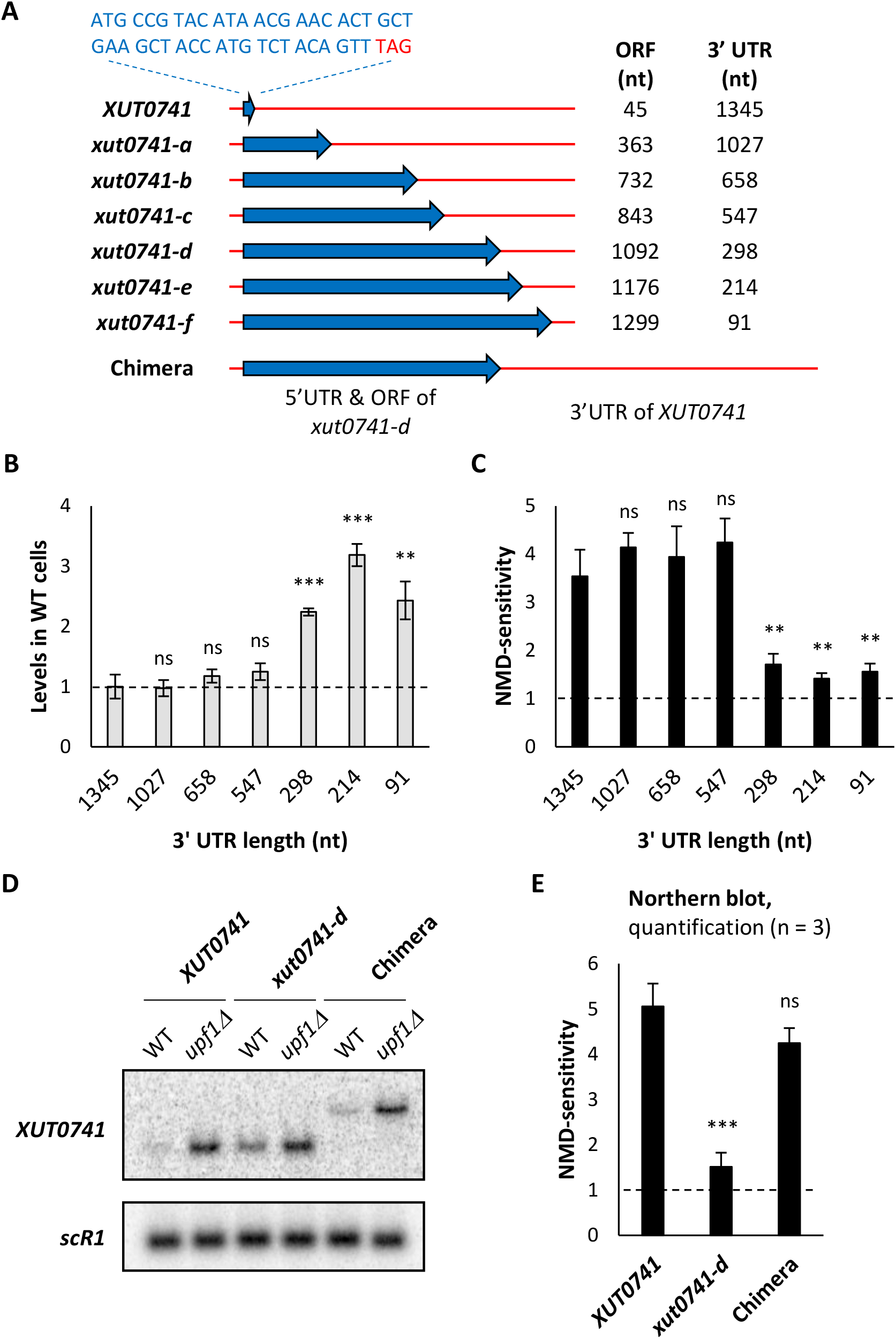
The NMD-sensitivity of *XUT0741* depends on its long 3’ UTR. **A.** Schematic representation of the native and mutant alleles of *XUT0741*. The transcript and the coding region are represented as a red line and a blue arrow, respectively. The sequence of the smORF in the native *XUT0741* is indicated. The length of the coding region and of the 3’UTR is shown beside each allele. The chimera construct was obtained by combining the 5’ UTR and coding region of the *xut0741-d* allele to the long 3’ UTR of the native *XUT0741*. **B.** WT and *upf1Δ* cells expressing the different alleles of *XUT0741* were grown to mid-log phase, at 30°C, in YPD medium. After total RNA extraction, the levels of each transcript were assessed by strand-specific RT-qPCR, and then normalized on *scR1*. The grey bars correspond to the levels of the different alleles of *XUT0741* in WT cells (native XUT set to 1, indicated by the dashed line). **C.** NMD-sensitivity of each allele of *XUT0741*, given by the ratio between the mean levels of each transcript in the *upf1* mutant and the WT strain (see Figure S6B). Mean and SD values were calculated from three independent biological replicates. ** *P* < 0.01; *** *P* < 0.001; ns, not significant upon t-test. The dashed line indicates a *upf1Δ*/WT ratio = 1 (*i.e*. no NMD-sensitivity). **D.** WT and *upf1Δ* cells expressing the native *XUT0741*, the *xut0741-d* allele and the chimera were grown as described above. Total RNA was extracted and analyzed by Northern blot. The different alleles of *XUT0741* and *scR1* (loading control) were detected using ^32^P-labelled AMO3581 and AMO1482 oligonucleotides, respectively. **E.** NMD-sensitivity of each allele, calculated from Northern blot signals in WT and *upf1Δ* cells (see Figure S6D). Mean and SD values were calculated from three independent biological replicates. *** *P* < 0.001; ns, not significant upon t-test. Dashed line: *upf1Δ*/WT ratio = 1.

Our data show that the abundance of the XUT in WT cells and its NMD-sensitivity remain unchanged in the three first mutants (Figure 6B-C; see also Figure S6B-C). However, when the 3’ UTR is shortened to 298 nt in the *xut0741-d* mutant (which is in the range of 3’ UTR size for NMD-insensitive XUTs; see Figure 5F), we observed a significant accumulation of the mutated transcript significant accumulation, correlating with a significant decrease of its sensitivity to NMD (Figure 6B-C; see also Figure S6B-C). Further shortening the 3’ UTR in mutants *–e* and *–f* did not aggravate these effects (Figure 6B-C). Note that the mutations introduced in *XUT0741* do not affect the NMD-sensitivity of another XUT (Figure S6C).

Thus, changing the length of the coding region relative to the 3’ UTR not only modifies the abundance of *XUT0741* in WT cells, but also its NMD-sensitivity. To discriminate whether the later depends on the length of the ORF or of the 3’ UTR, we constructed a chimera combining the extended ORF of ‘NMD-resistant’ *xut0741-d* with the long 3’ UTR of the native *XUT0741* (Figure 6A). The fate of this chimera was then analysed by Northern blot. As expected, the corresponding RNA was longer than the native XUT (Figure 6D). Importantly, the levels of the chimera in WT cells were undistinguishable from the levels of the native *XUT0741* (Figure 6D; see also Figure S6D), and in contrast to the *xut0741-d* mutant, the chimera and the native XUT display the same NMD-sensitivity (Figure 6E). We therefore conclude that the NMD-sensitivity of *XUT0741* is determined by its long 3’ UTR, which is consistent with the observation that the 3’ UTR for the translated NMD-sensitive XUTs is significantly longer (Figure 5F).

### Translation of an NMD-sensitive lncRNA produces a peptide in NMD-competent WT cells

All the observations described above contribute to the conclusion that translation occupies an important place in the metabolism of cytoplasmic lncRNAs. This led us to ask whether peptides could be produced as these lncRNAs are targeted to NMD (for simplicity, we will systematically use the term ‘peptide’ to refer to the product of the translation of a lncRNA, regardless its size).

Conceptually, the fact that NMD is triggered as translation terminates makes it possible for a peptide to be produced and released. To explore whether this could occur with yeast NMD-sensitive lncRNAs, we took advantage of the *xut0741-b* mutant (Figure 6A), which displays the same NMD-sensitivity as the native XUT (Figure 6C), but encodes a larger peptide and is easier to detect by Western blot. We decided to use this mutant as an artificial NMD-sensitive lncRNA reporter, following the insertion of a C-terminal 3FLAG tag (Figure 7A; see sequence in Supplementary File 1). We controlled that the insertion of the 3FLAG tag does not affect the NMD-sensitivity of the transcript (Figure 7B; see also Figure S7A). Importantly, despite the very low abundance of the transcript in WT cells, at the protein level we observed a clear band at the expected size by Western blotting, demonstrating that the encoded peptide is produced (Figure 7C, lane 3), its level increasing in the *upf1Δ* context (Figure 7C, lane 4). These results provide the proof-of-principle evidence that a peptide can be produced from an NMD-sensitive transcript in WT yeast cells.

**Figure 7.**
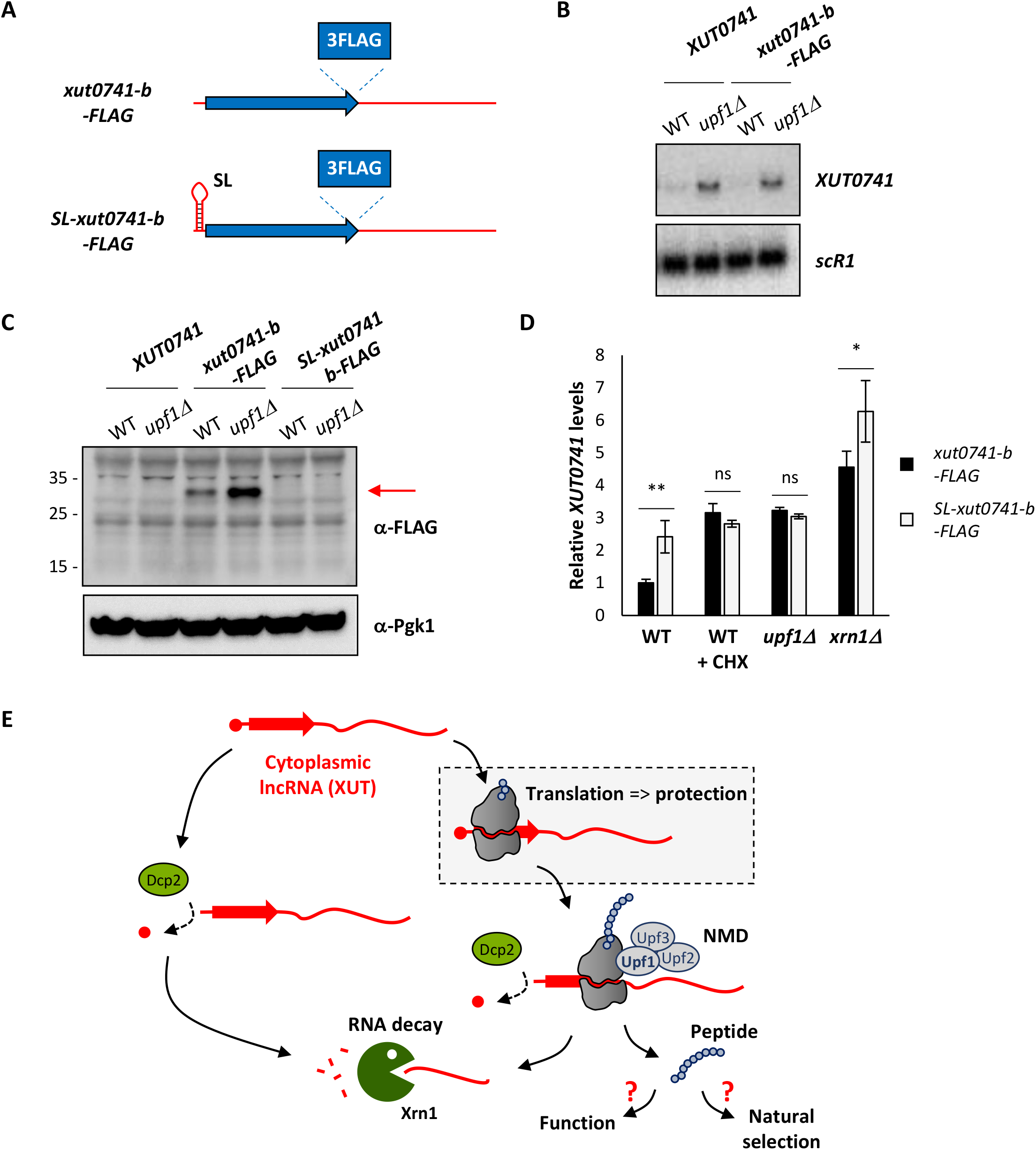
Detection of a translation product derived from NMD-sensitive XUT reporter in WT cells. **A.** Schematic representation of the tagged *xut0741-b* alleles, using the same color code as in Figure 6A. **B.** WT and *upf1Δ* cells expressing the native *XUT0741* or the *xut0741-b* allele fused to a C-terminal 3FLAG tag (*xut0741-b-FLAG*) were grown to mid-log phase, at 30°C, in YPD medium. Total RNA was extracted and analyzed by Northern blot. *XUT0741* and *scR1* were detected as described in Figure 1A. **C.** WT and *upf1Δ* cells expressing the native *XUT0741*, the *xut0741-b-FLAG* or the *SL-xut0741-b-FLAG* alleles were grown as above. Protein extracts (40 μg) were separated by poly-acrylamide gel electrophoresis and then transferred to a nitrocellulose membrane. The size of the protein ladder bands is indicated on the left of the panel. Pgk1 was used as a loading control. **D.** WT, *upf1Δ* and *xrn1Δ* cells expressing the *xut0741-b-FLAG* (black bars) or the *SL-xut0741-b-FLAG* (light grey bars) alleles were grown to mid-log phase, at 30°C, in YPD medium. For the WT strains, a sample of cells was also treated for 15 min with CHX (100 μg/ml, final concentration). After total RNA extraction, the levels of the corresponding transcript were assessed by strand-specific RT-qPCR, and then normalized on *scR1*. Mean and SD values were calculated from three independent biological replicates. The level of the *xut0741-b-FLAG* transcript in WT cells was set to 1. ** *P* < 0.01; * *P* < 0.05; ns, not significant upon t-test. **E.** Model. Translation of 5’ proximal smORF (red arrow) modulates the abundance of cytoplasmic lncRNAs. As they are translated, lncRNAs are protected from the degradation by the ribosomes. NMD is activated as translation terminates, leading to the degradation of the transcript. If not functional yet, the produced peptide could be exposed to the natural selection and possibly contributes to the progressive emergence of a *de novo* gene. The left part of the cartoon illustrates that in the absence of translation, lncRNAs are efficiently degraded, independently of NMD.

To gain further insight into the relationship between translation and NMD-sensitivity of XUTs, we designed a construct where the translation of our NMD-sensitive lncRNA reporter is blocked in *cis* by a short stem-loop (SL) element, known to inhibit translation initiation with a minimal effect on RNA decay (41,72). This SL was inserted into our reporter, upstream from the translation start site (Figure 7A; see sequence in Supplementary File 1). A Western blot showed that the production of the peptide is completely lost upon SL insertion, indicating that the transcript is no longer translated in this context (Figure 7C, lanes 5-6). At the RNA level, this loss of translation correlates with a dramatic decrease of the sensitivity of the XUT to CHX and to NMD (Figure 7D). Despite the effect of the SL sequence on Xrn1 was previously reported to be low (41,72), here we observed that it results in a significant accumulation of transcript in WT cells (Figure 7D; see also Figure S7B-C), probably impeding the exonuclease. However, this effect is partial as the transcript remains sensitive to Xrn1 (Figure 7D; see also Figure S7B-C).

The experimental detection of the translation product derived from our reporter upon epitope-tagging paves the way towards the characterization of the yeast peptidome using mass spectrometry (MS), having in mind that this has been reported to be technically challenging due to the limitations of the technique, especially for the detection of lowly-expressed microproteins (73). To date, we failed to detect the peptide derived from the native *XUT0741*, although we were able to unambiguously detect a synthetic, labelled version of it (see Figure S7D). We therefore postulate that the abundance of the native peptide within the total extract is not sufficient enough to be detected by MS.

Besides their low abundance, the amino acids composition of XUT-derived peptides might also impede their detection by MS. In fact, the composition of XUT-encoded peptides differs from the proteins encoded by mRNAs, with an over-representation of hydrophobic residues (such as phenylalanine, isoleucine and leucine), and an under-representation of lysine (see Figure S5E). We also noted the spectacular depletion of the negatively charged residues (aspartate and glutamate) in the peptides encoded by XUTs (see Figure S5E).

Notwithstanding the lack of robust MS data, our data show that translation of an NMD-sensitive lncRNA reporter gives rise to a peptide in WT cells, while the transcript is being efficiently targeted to NMD. Furthermore, our mechanistic analysis confirms that the CHX- and NMD-sensitivity of XUTs is indicative an active translation process.

## DISCUSSION

Since their discovery, lncRNAs have been considered as transcripts devoid of coding potential, thereby escaping translation. However, accumulating experimental evidence lead us to re-evaluate this assumption. In fact, lncRNAs co-purify with polysomes in different models, including yeast (15) and human cells (13,14,74). In addition, high-throughput sequencing of ribosome-bound fragments using Ribo-Seq or related approaches has uncovered smORFs within lncRNAs (11,12,16,17,74). Finally, several studies reported the identification of peptides resulting from the translation of smORFs carried on lncRNAs (16,19,21–23,74).

In yeast, lncRNAs expression is restricted by the extensive action of nuclear and cytoplasmic RNA decay machineries (75), including the 5’-exoribonuclease Xrn1 which degrades a family of cytoplasmic lncRNAs referred to as XUTs (37). We and others previously reported that most of them are targeted by the translation-dependent NMD pathway, suggesting that XUTs are translated and that translation controls their degradation (38,39).

Here we report several observations supporting this hypothesis. We showed that the majority of XUTs accumulate in WT cells treated with CHX or ANS (Figure 1), two drugs known to inhibit translation elongation but *via* different modes of action (53). Using Ribo-Seq we showed that a substantial fraction of XUTs are actually bound by ribosomes, and we identified actively translated smORFs which are mainly found in the 5’-proximal region of XUTs (Figure 4). Mechanistic analyses at the level of a candidate XUT showed that its sensitivity to NMD is determined by the length of the 3’ UTR downstream of the translated smORF (Figure 6). Finally, we showed that a detectable peptide is produced from an NMD-sensitive lncRNA reporter in WT cells as the transcript is targeted to NMD (Figure 7).

The fact that NMD-sensitive XUTs accumulate in the presence of a translation elongation inhibitor reinforces our model of a translation-dependent decay process. However, the underlying molecular mechanism appear to be more complex than anticipated, as the accumulation of XUTs observed upon CHX/ANS treatment cannot be solely explained by the inability of the cell to trigger NMD when translation is inhibited. Firstly, NMD-insensitive XUTs (which account for 30% of XUTs) also accumulate in presence of CHX or ANS (Figure 2A-B). Secondly, stress conditions associated with global translation initiation inhibition do not recapitulate the stabilization effect of the translation elongation inhibitors on XUTs (Figure 3B-C; see also Figure S3D). Thirdly, blocking elongating ribosomes with CHX interferes with the decapping of 35% of XUTs, which accumulate as capped RNAs in CHX-treated cells (Figure 2D), while most of the XUTs that accumulate upon NMD inactivation are decapped (Figure S2B-C). Together, these observations lead us to propose that while XUTs are translated, they would be protected by the elongating ribosomes sterically blocking the decay factors, independently of NMD. This model extends to lncRNAs the idea that translating ribosomes can protect mRNAs from the degradation (47,76).

Yet, several points remain to be clarified. Among them, the observation that the decapping of a fraction of XUTs is affected upon translation elongation inhibition raises the question of the difference between the XUTs that accumulate as capped or decapped in CHX-treated cells. Obviously, the fact that 65% of XUTs are efficiently decapped in these conditions rules out the possibility that CHX acts as a global inhibitor of decapping. Strikingly, most XUTs that accumulate as capped RNAs in CHX-treated WT cells are detected as translated by Ribo-Seq (Figure S5A-B). We found that XUTs abundance correlates with translation levels (Figure S5F) and resistance to decapping (Figure S5G). However, the functional relationship between XUTs translation and decapping appears to be more complex than anticipated. In fact, the Terminator-resistant and –sensitive subgroups of translated XUTs display the same 5’ UTR length (Figure S5C), as well as a similar ribosome occupancy along the 5’ region (Figure 5A and S5D). However, the Terminator-resistant XUTs display a significantly lower TE (Figure 5C). This suggests that when a XUT is translated with a low efficiency, it may remain bound by ribosomes for a longer period of time, which could negatively impact its decapping. Additional molecular analyses would be required to determine the mechanistic effect of TE on XUTs decapping. We also speculate that the dynamics of translation initiation could contribute to modulate the decapping of XUTs, possibly involving specific features or structures within the 5’ UTR that would interfere with the scanning process. Scanning the 5’ UTR until the first AUG codon does not depend on a fully assembled ribosome but on a 43S pre-initiation complex, which includes the 40S ribosome subunit, the initiator tRNA and several initiation factors (77). Technically, the classical Ribo-Seq approach we used detects translating ribosomes, but not the 43S pre-initiation complex. It would therefore be interesting to investigate the dynamics of scanning of the 5’ UTR of XUTs using 40S subunit profiling experiments (78,79), to determine whether and how this step affects the decapping and decay of XUTs.

The observation that most XUTs accumulate as decapped RNAs in *upf1Δ* cells was unexpected. On one hand, this shows that decapping remains efficient in the absence of NMD. On the other hand, how can one explain that XUTs accumulate and escape Xrn1 in this context, if they are decapped? Again, we could envisage a ribosome-mediated protection. In this regard, it has been shown that the ATPase activity of Upf1 is required for efficient ribosome release at the level of the stop codon; consequently, the inability to remove the terminating ribosome when this activity is lost impedes mRNA degradation by Xrn1, leading to the accumulation of 3’ mRNA decay fragments (64,80). However, this phenotype was only observed in the *upf1* ATPase mutant, but not in the *upf1Δ* null mutant. Additional works are therefore necessary to decipher the molecular mechanism leading to the stabilization of XUTs in the absence of functional NMD.

Our Ribo-Seq analysis allowed us to identify actively translated smORFs for 38% of annotated XUTs, including 510 NMD-sensitive XUTs and 123 NMD-insensitive XUTs (Figure 4D), considerably extending the repertoire of translated lncRNAs in yeast (15,39). Importantly, our data also indicate that NMD insensitivity does not imply lack of translation, and that the translational landscape of yeast lncRNAs extends beyond the scope of NMD. This is consistent with the observation that translation elongation inhibition also impacts the decay of most NMD-insensitive XUTs (Figure 2).

The number of smORFs/XUTs detected as translated depends on the stringency of the approach used to analyse the Ribo-Seq signals (Figure 4A), which is in line with the idea that lncRNA translation is transient and therefore more difficult to detect in comparison to canonical mRNAs translation (69). Besides the global low abundance of XUTs even in conditions where they are stabilized (NMD inactivation, CHX treatment), we imagine that the translation of many of their smORFs remains labile, probably reflecting the fact that they are rapidly and continuously evolving. This idea is supported by the idea that the vast majority of nORFs identified in yeast show no conservation (69). Furthermore, perhaps some constraints associated with canonical mRNA translation could be relaxed in the context of lncRNA translation as a strategy to maximize the potential for genetic novelty, which would be interesting from an evolutionary point of view. But the corollary is therefore the difficulty for us to detect such non-canonical translation events using pipelines that use the marks of canonical translation (*e.g*. use of an AUG initiator codon, predominance of one phase *vs* the two others). The field is therefore in need of dedicated approaches and computational tools to reveal the exhaustive landscape of lncRNA translation. Our data suggest that standard RNA-Seq analyses following a short treatment with translation elongation inhibitors are complementary to Ribo-Seq and could even represent an interesting alternative to it, simpler, cheaper and more sensitive, to reveal whether lowly abundant transcripts such as XUTs are translated or not.

Together with the observation that the NMD-sensitive XUTs display a longer 3’ UTR than the NMD-insensitive ones, the mechanistic analysis on the *XUT0741* candidate highlights the critical role of the 3’ UTR in determining the NMD-sensitivity of XUTs, as it is also the case for mRNAs (33,34). However, even in the mutant of *XUT0741* where the 3’ UTR is shortened to 91 nt, the NMD-sensitivity is not fully abolished (Figure S6B). One possible explanation is the existence of an alternative smORF, unaffected in our mutants. Supporting this hypothesis, we observed low Ribo-seq signals upstream from the detected smORF, overlapping the annotated TSS of *XUT0741* (Figure S6A). Interestingly, *XUT0741* TSS corresponds to the ‘G’ of an ‘ATG’ triplet, followed by 14 codons before the first in-frame stop codon (see sequences in Supplemental File 1). The production of multiple RNA isoforms from the same transcription unit is common in yeast (81), and we can imagine that any 5’-extended isoforms of *XUT0741* would encompass this ATG and therefore carry this alternative smORF. Additional mechanistic analyses combined to RNA isoform profiling would be required to confirm this hypothesis. Nonetheless, the complexity of the yeast transcriptome, with the existence of multiple RNA isoforms displaying different boundaries, might possibly explain the detection of several smORFs per XUT and should be kept in mind when investigating how the position of smORFs relative to its annotated extremities could impact the fate of a XUT.

Independently of the presence of a premature stop codon and of the length of the 3’ UTR, NMD-sensitive yeast mRNAs have been shown to be enriched non-optimal codons (34). Our data reveal that XUTs display an opposite pattern, the codon optimality score of the NMD-sensitive XUTs being significantly higher in comparison to the NMD-insensitive XUTs (Figure 5E). More globally, the codon optimality score of XUTs is significantly lower compared to mRNAs (Figure 5D), which would be consistent with their high instability (70). Together with the different composition in amino acids of XUT-derived peptides in comparison to proteins (Figure S7E), this also indicates that the primary properties of the smORFs of XUTs are distinct from the canonical ORFs of yeast, possibly reflecting that they are in a very early stage of selection, if only engaged in this process.

One important finding of our work is that the translation of an NMD-sensitive lncRNA reporter gives rise to a peptide detectable in a WT context, where NMD is functional. From a conceptual point of view, the idea that a peptide can be produced from an NMD substrate is plausible, since NMD is activated as translation terminates at the level of a ‘normal’ stop codon (this is the position of this codon within the transcript which is sensed as ‘abnormal’). However, the fate of this peptide has not been characterized in detail so far and remains largely obscure. On one side, a study in yeast proposed that Upf1 stimulates the proteasome-dependent degradation of the truncated translation product derived from an NMD-sensitive mRNA carrying nonsense mutation (82), consistent with the classical view that such products might be deleterious for the cell and should be eliminated. In contrast, a study in mammalian cells revealed that the pioneer round of the translation which targets mRNAs with premature stop codons to NMD can produce antigenic peptides for the MHC class I pathway (83). In this context, the observation we made here using our tagged NMD-sensitive reporter provides the proof-of-principle that translation of an NMD-sensitive transcript can give rise to a peptide which can exist in the cell, even if the transcript it originates from is targeted for degradation. This first observation paves the way towards the future characterization of the yeast peptidome using mass spectrometry, searching for native peptides derived from the translation of XUTs, keeping in mind the actual technical limitations of this approach for the identification of lowly-expressed and short peptides (18,73).

Overall, our data lead us to propose that translation of 5’-proximal smORFs is a general feature of the cytoplasmic lncRNAs, modulating their cellular abundance (Figure 7E). While they are translated, the presence of elongating ribosomes would protect them from the decay factors. Then, as translation of these smORFs terminates far away from the poly(A) tail, the NMD factors would be recruited to the terminating ribosome, triggering NMD. At the same time, the peptides that have been produced could be exposed to natural selection, and if beneficial, they could be selected and ultimately lead to *de novo* protein-coding genes.

*De novo* gene birth has been associated with adaptation to environmental stress (84), and NMD is known to be repressed under a variety of stress conditions (85,86). It is therefore tempting to speculate that despite the fact that the cell has evolved efficient pathways to degrade lncRNAs and restrict their expression, these pathways can be down-regulated under certain conditions (*e.g*. stress) in order to sample the peptide potential hosted in these lncRNAs.

An important corollary of our model is that lncRNA-derived peptides are unlikely to be immediately functional. Consequently, their loss is not expected to confer a phenotype. However, their overexpression might confer a selective advantage. This thought highlights the importance of addressing the question of the functionality of lncRNAs by considering approaches based on gain-of-function (87,88). However, we do not exclude the possibility that some lncRNA-derived peptides could be functional and confer a growth phenotype when they production is abolished, as shown for mutants of the AUG codon of three yeast nORFs (89).

In conclusion, our work is coherent with the idea that translation plays a major role in the post-transcriptional metabolism of cytoplasmic lncRNAs, and that their definition as ‘non-coding’ is probably not an appropriate description of their actual status. Rather, they might be viewed as transcripts oscillating between the ‘coding’ and ‘non-coding’ worlds, sources of potential of genetic novelty *via* the production of novel peptides, which if beneficial for the cell, might be selected to give rise to novel protein-coding genes.

## FUNDING

This work has benefited from the ANR “DNA-life” (ANR-15-CE12-0007) grant and the ERC “DARK” consolidator grant allocated to A. M. The ICGex NGS platform of the Institut Curie is supported by the grants ANR-10-EQPX-03 (Equipex) and ANR-10-INBS-09-08 (France Génomique Consortium) from the Agence Nationale de la Recherche (“Investissements d’Avenir” program), by the ITMO-Cancer Aviesan (Plan Cancer III) and by the SiRIC-Curie program (SiRIC Grant INCa-DGOS-465 and INCa-DGOSInserm_12554). S. A. has been supported by a PhD fellowship from PSL university and by the Fondation pour la Recherche Médicale (FRM).

## ACKNOWLEDGEMENTS

We would like to thank Aaron Wacholder and Anne-Ruxandra Carvunis for sharing unpublished results and fruitful discussions. We are also grateful to all members of our labs for discussions and to our colleague Michael Schertzer for his careful reading of this manuscript and English revisions. High-throughput sequencing was performed by the ICGex NGS platform of the Institut Curie. Data management, quality control and primary analysis were performed by the Bioinformatics platform of the Institut Curie.

## AUTHOR CONTRIBUTIONS

M.W., I.H., O.N. & A.M. designed experiments. S.A., N.V., I.H., D.C. & M.W. performed experiments. U.S., C.P. & A.L. performed bioinformatics analyses. S.A., U.S., I.H., D.C., C.P., A.L. & M.W. analysed the data. M.W. & A.M. designed the project. M.W. & A.M. supervised the project. M.W. wrote the manuscript, with input from all authors. A.M. acquired funding.

## CONFLICT OF INTEREST

The other authors declare that they have no competing interests.

## Notes

### Competing Interest Statement

The authors have declared no competing interest.

### Summary of Updates

Figure 3 and 7 have been revised, and a novel figure (Figure 5) has been included.

