## Supplemental Figures 1-8 for "Translation is a key determinant controlling the fate of cytoplasmic long non-coding RNAs"

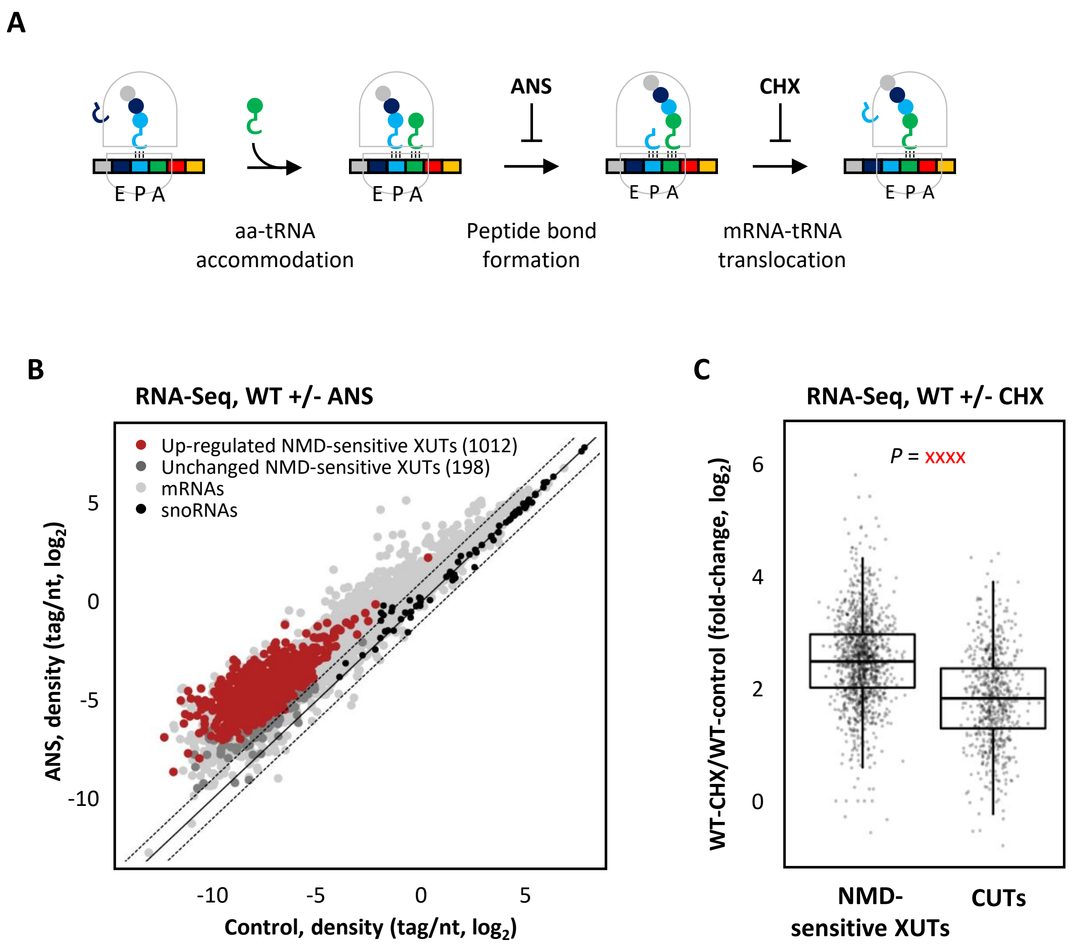


**Figure S1. NMD-sensitive lncRNAs accumulate upon translation inhibition.**

**A.** Schematic representation of the eukaryotic translation elongation cycle. The steps specifically inhibited by CHX and ANS are highlighted. The codons on the mRNA, the tRNAs and the amino acids (aa) are represented as rectangles, loops and circles, respectively (a color code is used to show the codon/tRNA/aa correspondence). The E, P and A sites of the ribosome are indicated. **B.** Total RNA-Seq was performed using total RNA extracted from exponentially growing WT (YAM1) cells (grown as above) treated for 30 minutes with ANS (100 µg/ml, final concentration) or with an equal volume of DMSO (control). The scatter plot shows the RNA-Seq signals (tag densities, $log_2$ scale) for the NMD-sensitive XUTs, mRNAs (light grey dots) and snoRNAs (black dots) in ANS-treated and control WT cells. The significantly up-regulated (ANS/control fold-change >2, *P*-value <0.05) and unaffected NMD-sensitive XUTs are represented as red and dark grey dots, respectively. **C.** Sensitivity of NMD-sensitive XUTs and CUTs to CHX. The box-plot shows the global sensitivity to CHX of NMD-sensitive XUTs and 'strict' CUTs, in WT cells (CHX/control ratio of RNA-Seq signals). The 'strict' CUTs correspond to a subgroup of CUTs (621) that do not overlap XUTs.

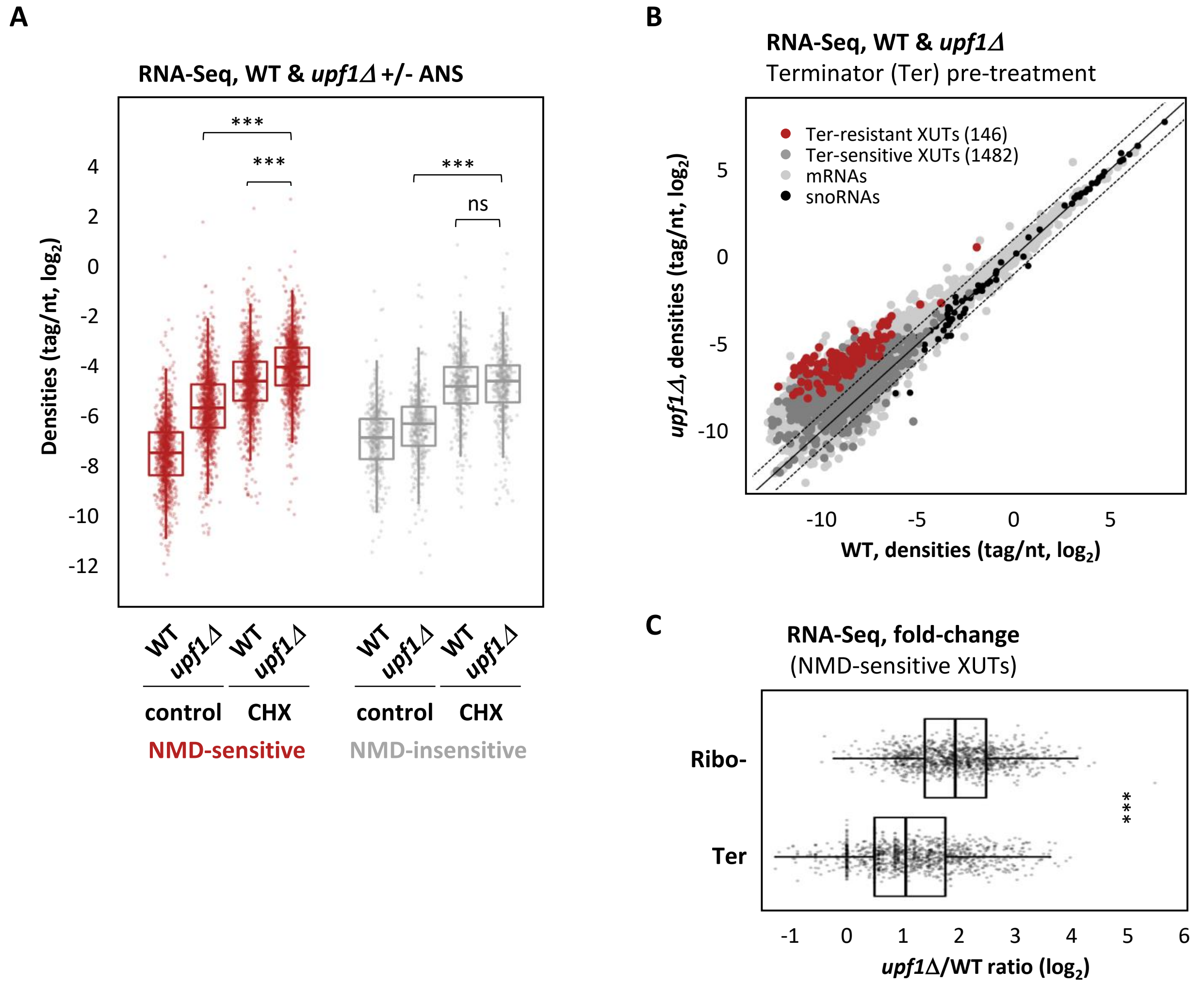


**Figure S2. Translation also impacts XUTs independently of NMD.**

**A.** Total RNA-Seq was performed in WT (YAM1) and *upf1Δ* (YAM202) cells treated for 30 minutes with ANS (100 µg/ml, final concentration) or an equal volume of DMSO. The box-plot shows the densities (tag/not, $log_2$) computed for the NMD-sensitive and NMD-insensitive XUTs. *** *P*-value < 0.001; ns, not significant upon two-sided Wilcoxon rank-sum test (adjusted for multiple testing with the Benjamini–Hochberg procedure). **B.** Total RNA-Seq was performed using total RNA extracts from WT (YAM1) and *upf1Δ* (YAM202) cells, including a treatment with the Terminator 5'-phosphate-dependent exonuclease (which digests decapped RNAs) before the preparation of the libraries. The data are presented as a scatter plot showing the RNA-Seq signals (tag densities, $log_2$ scale) for the NMD-sensitive XUTs, mRNAs (light grey) dots and snoRNAs (black dots). The red dots represent the NMD-sensitive XUTs that are still detected as significantly up-regulated in *upf1Δ* cells (*upf1Δ*/WT fold-change >2, *P*-value <0.05) upon Terminator treatment. The other XUTs (Terminator-sensitive) are represented as dark grey dots. **C.** Box-plot of the *upf1Δ*/WT fold-change for the NMD-sensitive XUTs computed using RNA-Seq data obtained from libraries prepared using total RNA extracts submitted to rRNA depletion (Ribo-) or Terminator digestion (Terminator).

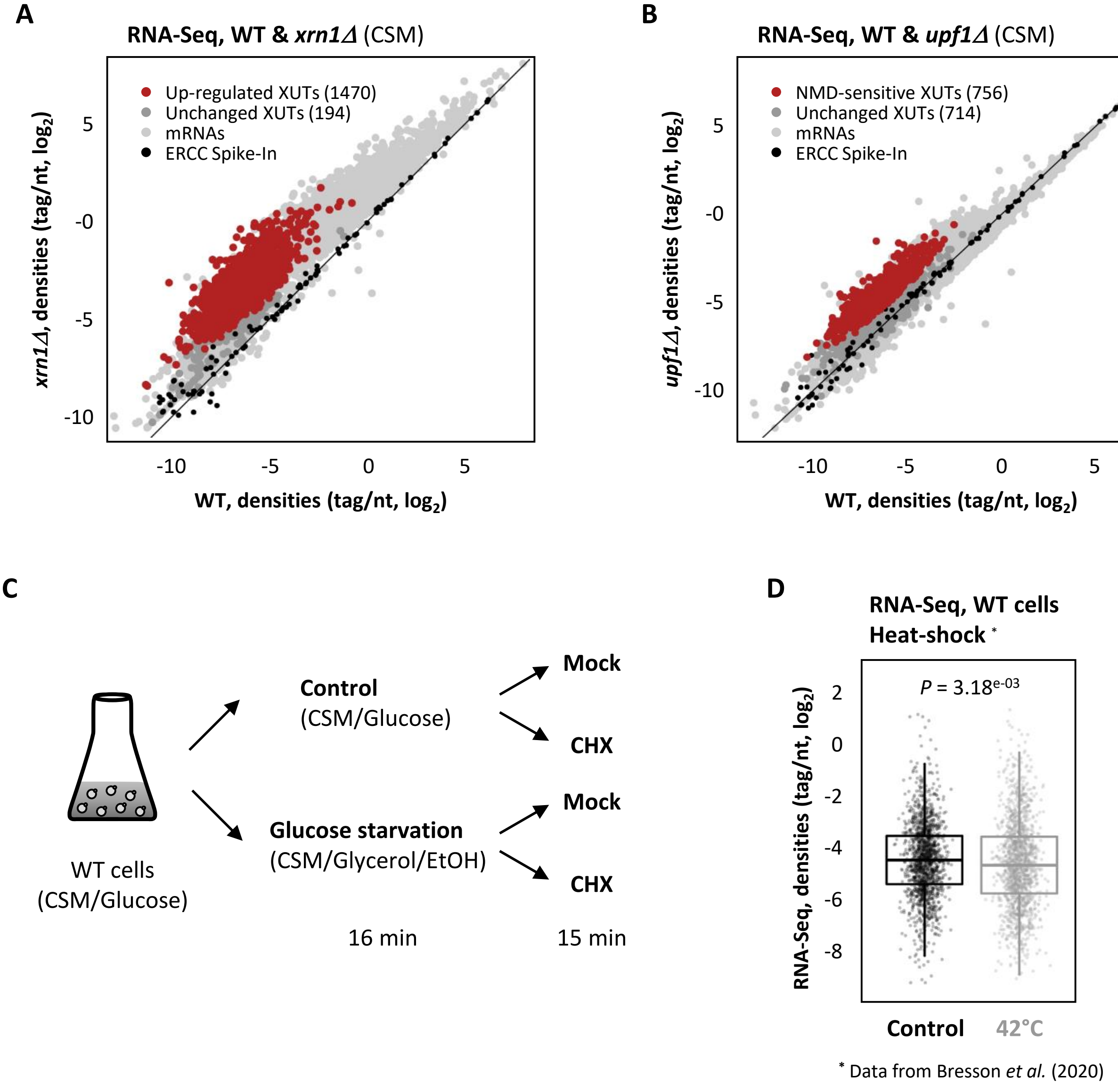


**Figure S3. XUTs levels remain unaffected upon translation initiation inhibition.**
**A.** XUTs landscape in CSM medium. Total RNA-Seq was performed in WT (YAM1) and *xrn1Δ* (YAM6) cells grown to mid-log phase in Complete Synthetic Medium (CSM). Densities (tag/not, $log_2$) were computed for XUTs, mRNAs (light grey dots) and snoRNAs (black dots). The 1470 XUTs up-regulated in the *xrn1* mutant (*xrn1Δ*/WT fold-change >2, *P*-value <0.05) are highlighted in red. The dark grey dots correspond to the other XUTs, the expression of which is not significantly affected. **B.** Landscape of NMD-sensitive XUTs in CSM medium. Same as above, using WT (YAM1) and *upf1Δ* (YAM202) cells grown in CSM. The red dots represent the 756 XUTs defined as NMD-sensitive in this condition (*upf1Δ*/WT fold-change >2, *P*-value <0.05). **C.** Experimental scheme. WT (YAM1) cells were grown to mid-log phase in CSM with glucose as carbon source, and then shifted for 16 min in CSM where glucose was replaced by glycerol and ethanol (glucose starvation). In parallel, control cells were maintained for the same time in glucose-containing CSM. CHX (100 µg/ml final concentration) or an equal volume of DMSO (Mock) was then added to each sample. Cells were harvested after 15 min of treatment; then total RNA was extracted. Note that the CSM medium used here is different from the rich medium (YPD) that was originally used to annotate XUTs, so that we had to re-define the XUTs landscape in CSM (see above). **D.** Analysis of published RNA-Seq data obtained in WT cells grown in CSM and then shifted for 16 min at 42°C (47). Densities (tag/nt, $log_2$) were computed for the 1470 XUTs expressed in CSM (see panel A). The indicated *P*-value was obtained upon two-sided Wilcoxon rank-sum test.

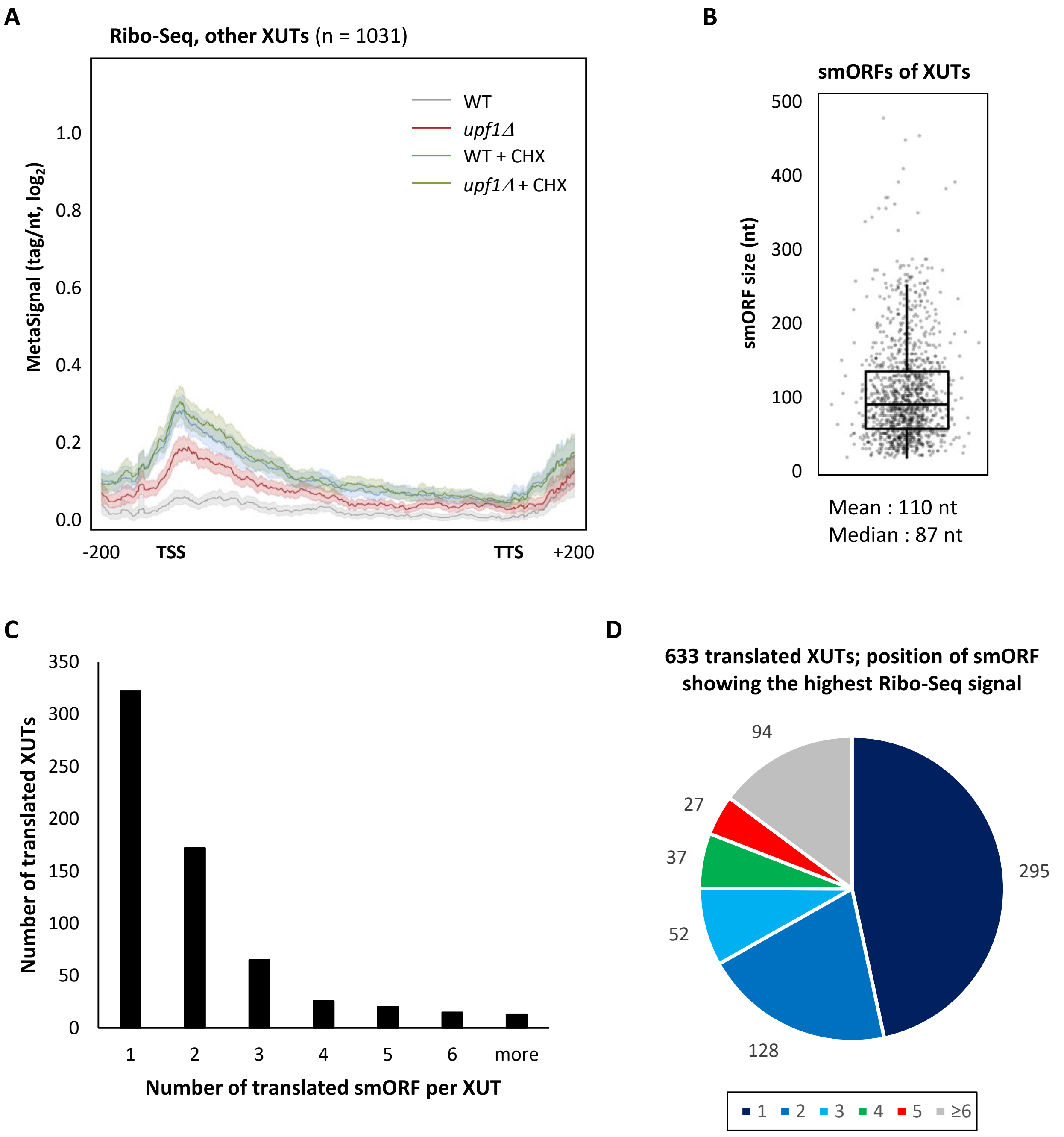


**Figure S4. Translational landscape of XUTs.**

**A.** Metagene of Ribo-Seq signals along the 1031 XUTs that were not detected as translated upon analysis using the Ribotricer method, separating the different conditions (*i.e.* XUTs excluded from list 2). For each condition, the densities (tag/nt, $\log_2$) along the XUTs +/- 200 nt were piled up, then the average signal was plotted. The shading surrounding each line denotes the 95% confidence interval. **B.** Box-plot representation of the size of the 1270 translated smORFs of XUTs (list 2). The mean and median values are indicated. **C.** Histogram showing the number of translated smORFs per XUTs (for the 1270 smORFs and 633 XUTs of list 2). **D.** Pie chart showing for the 633 translated XUTs (list 2) the position of the smORF with the highest Ribo-Seq signal relative to all the smORFs predicted across the XUT sequence (≥ 5 codons, starting with an AUG).

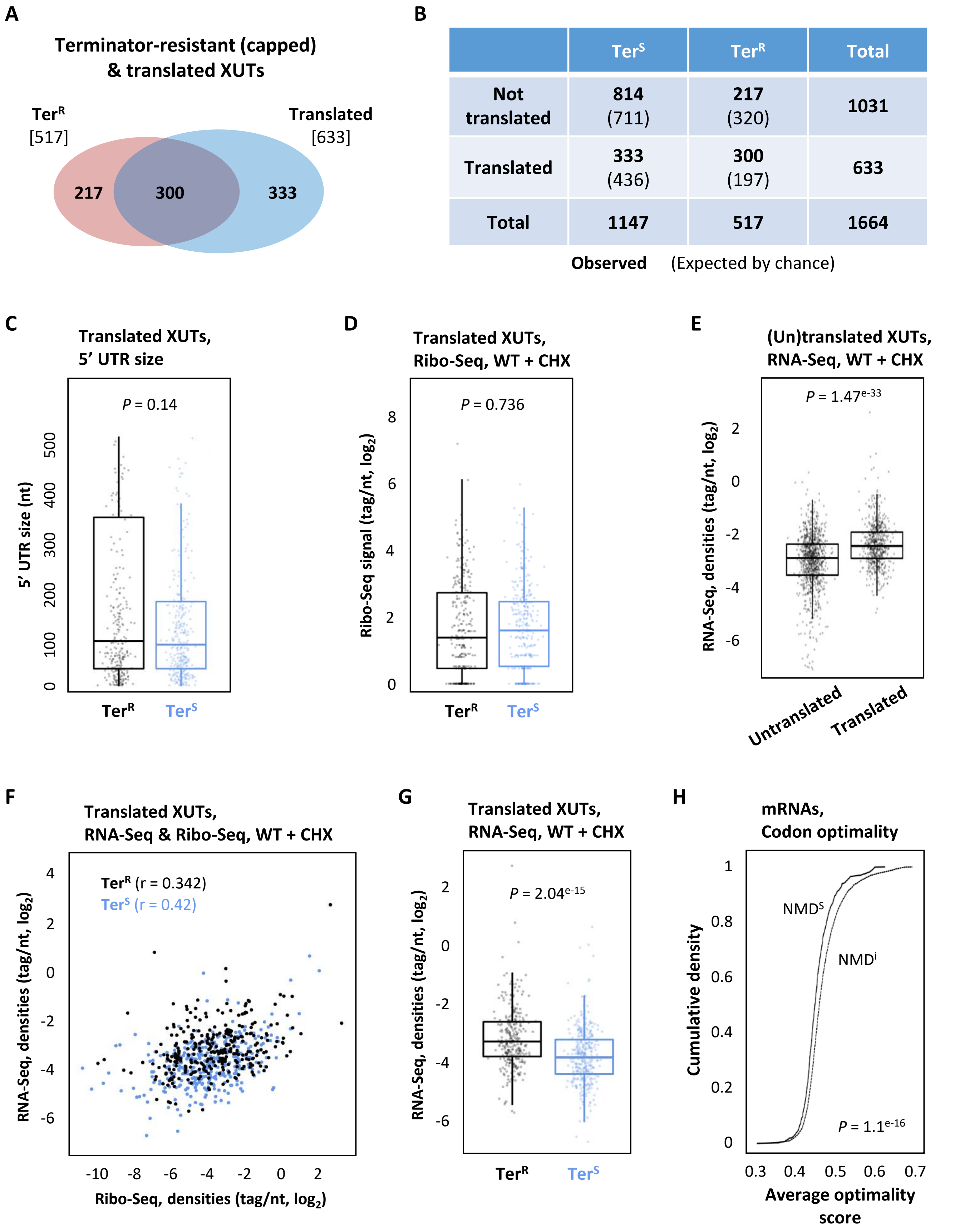


**Figure S5. Legends on next page**

**Figure S5. Features of translated XUTs**

**A.** Venn diagram showing the number of XUTs that accumulate as Terminator-resistant ($Ter^R$) RNAs in CHX-treated cells and that are detected as translated by Ribo-Seq. **B.** Contingency table displaying the number of XUTs detected as translated or not by Ribo-Seq and the number of XUTs that accumulate as Terminator-resistant ($Ter^R$) or -sensitive ($Ter^S$) RNAs in CHX-treated WT cells. Observed numbers are in bold. Numbers expected by chance are indicated in bracket. The *P*-value obtained upon chi-square test of independence is indicated. **C.** Box-plot showing the size of the 5' UTR for the 300 Terminator-resistant ($Ter^R$, black) and the 333 Terminator-sensitive ($Ter^S$, blue) translated XUTs. The smORF with the highest Ribo-Seq signal was considered for XUTs with several annotated smORFs. The *P*-value obtained upon two-sided Wilcoxon rank-sum test is indicated. **D.** Ribo-Seq signals (tag/nt, $log_2$) along the TSS to TSS+50 region for the Terminator-resistant and Terminator-sensitive translated XUTs, in CHX-treated cells. The *P*-value was obtained upon two-sided Wilcoxon rank-sum test. **E.** RNA-Seq signals in CHX-treated WT cells for the XUTs detected as translated (633) or not (1031). **F.** Scatter plot showing the Ribo-Seq (TSS to TSS+50) and RNA-Seq signals (tag/nt, $log_2$ scale) for the Terminator-resistant ($Ter^R$, black dots) and the Terminator-sensitive ($Ter^S$, blue dots) translated XUTs. The Pearson correlation coefficient (r) is indicated for each subgroup of XUTs. **G.** RNA-Seq signals (tag/nt, $log_2$) for the Terminator-resistant and Terminator-sensitive translated XUTs, in CHX-treated cells. The indicated *P*-value was obtained upon two-sided Wilcoxon rank-sum test. **H.** Average codon optimality score for the NMD-sensitive ($NMD^S$, full line, n = 833) and NMD-insensitive ($NMD^i$, dashed line, n = 4667) mRNAs, shown as a cumulative frequency plot. The indicated *P*-value was obtained upon Kolmogorov–Smirnov test.

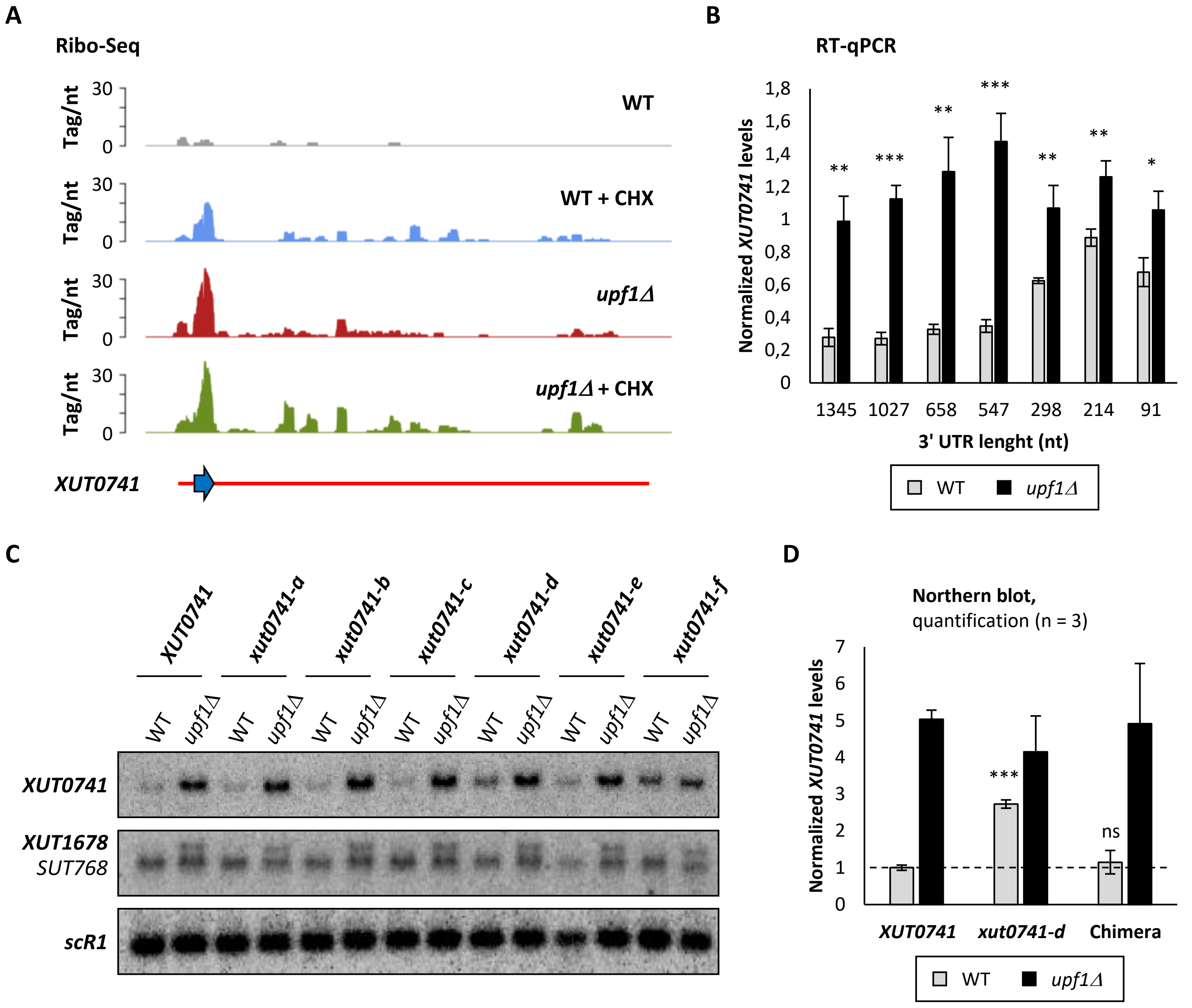


**Figure S6. The NMD-sensitivity of *XUT0741* depends on its long 3' UTR.**

**A.** Snapshot of Ribo-Seq signals across *XUT0741* in WT and *upf1Δ* cells, with or without CHX treatment. For each condition, the signals (tag/nt) obtained for the two biological replicates were added. *XUT0741* is depicted as a red line. The blue arrow represents the single smORF detected as actively translated in our analysis. **B.** WT and *upf1Δ* cells expressing the different alleles of *XUT0741* (see Figure 6A) were grown to mid-log phase at 30°C in YPD medium. After total RNA extraction, the levels of each transcript were assessed by strand-specific RT-qPCR, and then normalized on *scR1*. The black and grey bars represent the mean values +/- SD in WT (grey) and *upf1Δ* (black) cells, calculated from three independent biological replicates. $^{*}$ $P < 0.05$; $^{**}$ $P < 0.01$; $^{***}$ $P < 0.001$ upon t-test. **C.** The same RNAs as above were analyzed by Northern blot. *XUT0741*, *XUT1678* (and the overlapping *SUT768*) and *scR1* (loading control) were detected using $^{32}$P-labelled AMO1762, AMO1595 and AMO1482 oligonucleotides, respectively. **D.** WT and *upf1Δ* cells expressing the native *XUT0741*, the *xut0741-d* allele and the chimera were grown as described above. Total RNA was extracted and analyzed by Northern blot. The different alleles of *XUT0741* and *scR1* (loading control) were detected by Northern blot using $^{32}$P-labelled AMO3581 and AMO1482 oligonucleotides, respectively. Mean and SD values were calculated from three independent biological replicates. The average level of the native *XUT0741* in WT cells was set to 1. $^{***}$ $P < 0.001$; ns, not significant upon t-test.

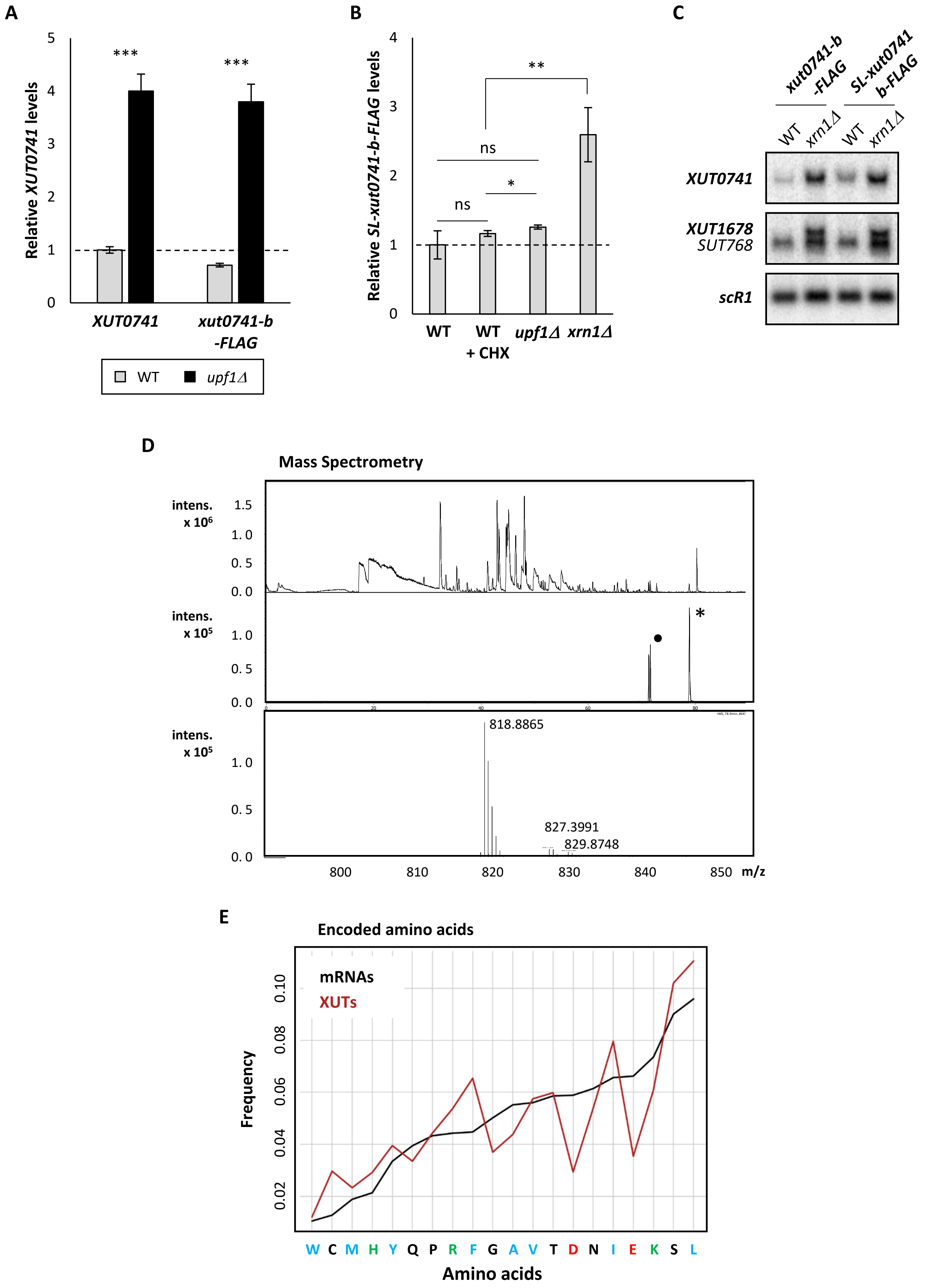


Figure S7. Legends on next page

**Figure S7. Detection of a translation product derived from NMD-sensitive XUT reporter in WT cells.**

**A.** WT and *upf1Δ* cells expressing the native *XUT0741* or the *xut0741-b* allele fused to a C-terminal 3FLAG tag (*xut0741-b-FLAG*) were grown to mid-log phase, at 30°C, in YPD medium. Total RNA was extracted and analyzed by strand-specific RT-qPCR. Levels of *XUT0741* transcripts were normalized on *scR1*. Mean and SD values were calculated from three independent biological replicates. *** $P < 0.001$ upon t-test. Dashed line: WT levels of *XUT0741* set as 1. **B.** WT, *upf1Δ* and *xrn1Δ* cells expressing the *SL-xut0741-b-FLAG* allele were grown as above. For the WT strain, a sample of cells was also treated for 15 min with CHX (100 µg/ml, final concentration). After total RNA extraction, the levels of the *SL-xut0741-b-FLAG* transcript were assessed by strand-specific RT-qPCR, normalized on *scR1* and set as 1 for the untreated WT condition (indicated by the dashed line). Mean and SD values were calculated from three independent biological replicates. ** $P < 0.01$; * $P < 0.05$; ns, not significant upon t-test. **C.** WT and *xrn1Δ* cells expressing the *xut0741-b-FLAG* or the *SL-xut0741-b-FLAG* alleles were grown as above. Total RNA was extracted and analysed by Northern-blot. *XUT0741*, *XUT1678* (and the overlapping *SUT768*) and *scR1* (loading control) were as described in Figure S6B. **D.** Mass spectrometry analysis of the 1-10 kDa gel fraction from *upf1Δ* (YAM202) cells grown in the presence of MG132 proteasome inhibitor, spiked with the *XUT0741* heavy synthetic peptide. Base peak chromatogram (upper panel) and MS extracted-ions chromatogram (XIC) of the *XUT0741* labelled peptide (m/z = 818.88) marked with an asterisk (middle panel) are showed. Mass spectrum (lower panel) acquired at the retention time of the heavy synthetic peptide show the absence of peak at lower mass (m/z = 815.4) corresponding to its endogenous counterpart. Chromatographic peak marked with bold circle corresponds to ion interference. **E.** Frequency of amino acids encoded by the ORFs of mRNAs (black curve) or the smORFs of XUTs (red curve). The smORF with the highest Ribo-Seq signal was considered for XUTs with several annotated smORFs. Hydrophobic, positively charged and negatively charged residues are indicated in blue, green and red respectively. The other residues are in black.

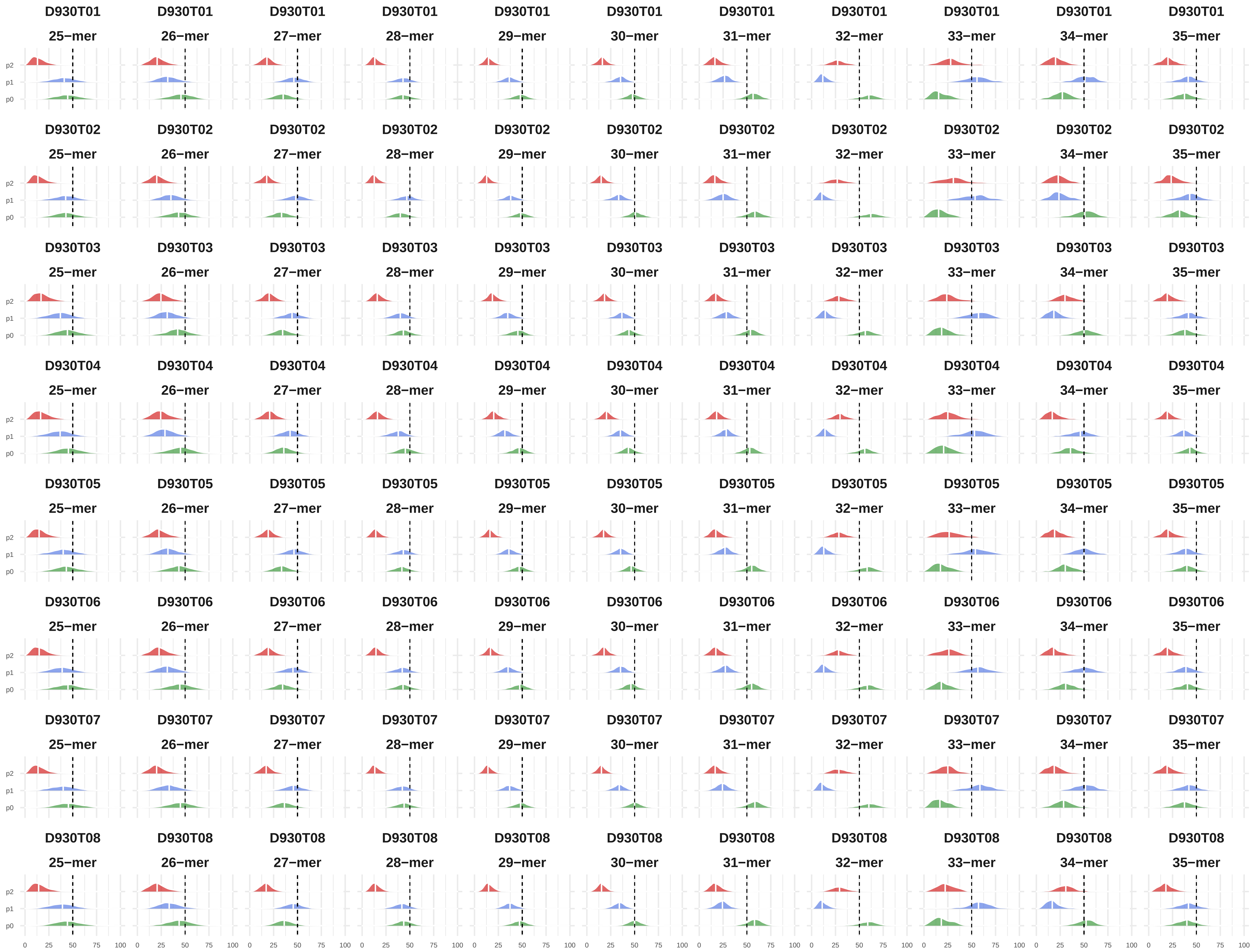
D930T01 25−mer
D930T01 26−mer
D930T01 27−mer
D930T01 28−mer
D930T01 29−mer
D930T01 30−mer
D930T01 31−mer
D930T01 32−mer
D930T01 33−mer
D930T01 34−mer
D930T01 35−mer
D930T02 25−mer
D930T02 26−mer
D930T02 27−mer
D930T02 28−mer
D930T02 29−mer
D930T02 30−mer
D930T02 31−mer
D930T02 32−mer
D930T02 33−mer
D930T02 34−mer
D930T02 35−mer
D930T03 25−mer
D930T03 26−mer
D930T03 27−mer
D930T03 28−mer
D930T03 29−mer
D930T03 30−mer
D930T03 31−mer
D930T03 32−mer
D930T03 33−mer
D930T03 34−mer
D930T03 35−mer
D930T04 25−mer
D930T04 26−mer
D930T04 27−mer
D930T04 28−mer
D930T04 29−mer
D930T04 30−mer
D930T04 31−mer
D930T04 32−mer
D930T04 33−mer
D930T04 34−mer
D930T04 35−mer
D930T05 25−mer
D930T05 26−mer
D930T05 27−mer
D930T05 28−mer
D930T05 29−mer
D930T05 30−mer
D930T05 31−mer
D930T05 32−mer
D930T05 33−mer
D930T05 34−mer
D930T05 35−mer
D930T06 25−mer
D930T06 26−mer
D930T06 27−mer
D930T06 28−mer
D930T06 29−mer
D930T06 30−mer
D930T06 31−mer
D930T06 32−mer
D930T06 33−mer
D930T06 34−mer
D930T06 35−mer
D930T07 25−mer
D930T07 26−mer
D930T07 27−mer
D930T07 28−mer
D930T07 29−mer
D930T07 30−mer
D930T07 31−mer
D930T07 32−mer
D930T07 33−mer
D930T07 34−mer
D930T07 35−mer
D930T08 25−mer
D930T08 26−mer
D930T08 27−mer
D930T08 28−mer
D930T08 29−mer
D930T08 30−mer
D930T08 31−mer
D930T08 32−mer
D930T08 33−mer
D930T08 34−mer
D930T08 35−mer
p2
p1
p0
0 25 50 75 100

**Figure S8. Phasing of Ribo-Seq data.**
For each dataset, the P-site of the different k-mers (25-mers to 35-mers) was predicted with RiboWaltz (78). As a quality control, for each k-mer, we calculated for the protein-coding genes the fraction of reads that are in-frame with the expected ORF (mentioned as P0). D930T01-02 : WT – native conditions, replicates 1-2 ; D930T03-04 : WT – CHX, replicates 1-2 ; D930T05-06 : *upf1Δ* – native conditions, replicates 1-2 ; D930T07-08 : *upf1Δ* – CHX, replicates 1-2.
